# The immune and genetic heterogeneity of hepatocellular carcinoma, with a focus on multifocal disease

**DOI:** 10.1101/2025.10.01.678727

**Authors:** Abigail Connor, Leon Chang, Emma West, Karen Scott, Fay Ismail, Alyn Cratchley, Darren Newton, Adel Samson

**Affiliations:** Leeds Institute of Medical Research at St. James’s, University of Leeds, Leeds, United Kingdom; Leeds Teaching Hospitals NHS Trust, Leeds, United Kingdom

**Keywords:** Hepatocellular carcinoma, tumour immunology, genomic profiling, multifocal cancer

## Abstract

Understanding the genetic and immune heterogeneity in hepatocellular carcinoma (HCC) is crucial as treatments advance towards personalisation. This study characterises the genetic and immune heterogeneity within a population of 56 tumours from 28 patients, 10 of whom have multifocal disease, obtained from patients undergoing surgery at the Leeds Teaching Hospital NHS Trust. These samples represent the most common aetiological groups found in the UK: alcohol-related liver disease, non-alcoholic fatty liver disease and hepatitis C. Whole exome sequencing was performed on DNA from these tumour alongside matched background liver samples to determine tumour-specific variants. Tissue sections were stained for both proliferation and immune cell markers. Tumours across aetiological groups showed significant levels of variation, with high inter-individual variation. Analyses of multifocal tumours revealed significant discordance in genetic and immune cell profiles, both between multi-centric primary and metastatic tumours. This work emphasises the genetic and immuneheterogeneity in HCC across HCC subtypes, between and within individuals, highlighting mechansisms for therapeutic resistance and the need for personalised medicine.

## Introduction

Hepatocellular carcinoma (HCC) is the third-largest cause of cancer-related deaths worldwide, responsible for over 800,000 deaths annually (Sung et al., 2021). Global incidence is projected to increase further, driven by the rising prevalence of underlying risk factor; most commonly by alcohol-related liver disease (ARLD), non-alcoholic fatty liver disease (NAFLD) and hepatitis B/C virus (HBV/HCV) infections (Fitzmaurice et al., 2017).

HCCs are not defined by a singular genetic event but rather an enrichment of mutations, occurring in specific disease-associated geness. A recent analysis of The Cancer Genome Atlas (TCGA) Liver Hepatocellular Carcinoma (LIHC) dataset revealed that the variants are most often misense single nucleotide polymorphisms, with the *Titin* (*TTN)* gene containing one or more variants in 24% of tumours (Zheng et al., 2024). Compared to other tumour types, HCC has an intermediate tumour mutation burden (TMB; Chalmers et al., 2017), and literature suggests this is often variable: Wong et al. (2021) reported a range between 0 and 3.5 mutations per megabase in sequenced tumours. TMB has been linked to immune checkpoint blockade (ICB) response in a meta-analysis of solid cancers, where a 48% reduced risk of disease progression was observed in patients with a ‘high’ TMB following ICB treatment (Kim et al., 2019).

One mechanistic hypothesis underlying this observation is that a higher TMB results in a higher frequency of tumour-specific mutations which, in turn, increases the neoepitope load of a tumour. A higher frequency of non-self-proteins presented on the surface of cancer cells creates more opportunities for T cell recognition and subsequent tumour killing. As current immunotherapies work, in part, by increasing the activity of T cells, efficacy may be more limited in tumours with a low TMB.

Consistent with other cancer types, the frequency of tumour-infiltrating lymphocytes is associated with overall survival (OS) in HCC (Ding et al., 2018). Whilst the FoxP3+ subset is instead linked with poorer survival (Fu et al., 2007). These observations support the concept that the efficacy of immunotherapy is at least partially dependant on a pre-existing anti-tumour immune response, as seen in non-small cell lung cancer patients (Xagara et al., 2024).

Distinct risk factors have been linked to the aforementioned aetiological groups underlying HCC, but how these influence genetic and immune cell variation is poorly characterised. Samples in South-East Asia have often been reported to contain the TP53 R249S hotspot mutation, which is linked to exposure to aflatoxin metabolites (Galy et al., 2011). All aetiologies have been associated with immune cell exhaustion (Llovet et al., 2022), whilst responses to immunotherapy have been reported to vary between groups (Pfister et al., 2021). Characterising the extent of variation in genetic alterations and immune cell frequencies between HCC in this current cohort offers key insights into likely reasons for resistance to current immunotherapy treatments, including the complexities of multifocal disease. Advancements in our understanding of tumour biology has allowed the stratification of treatment approaches in certain cancer types. However, at the time of this study, patients with advanced HCC are most often treated with a combination of Atezolizumab and Bevacizumab, despite favourable responses observed in only a minority of cases.

In this study, we characterised a set of 56 tumours from 28 patients undergoing liver transplant surgery, including 10 with multifocal disease. Our analyses further the understanding of the immunogenetic landscape of HCC and provides an opportunity to compare the tumour immune landscape of multicentric HCC. Characterising the extent of variation in genetic alterations and immune cell frequencies between HCCs in this cohort offers key insights into likely reasons for resistance to current immunotherapy treatments, including the complexities of multifocal disease, providing evidence to aid the rational development of novel threrapeutics.

## Methods

### Statistics and data visualisation

P values were calculated using one-way ANOVA, paired T tests or Pearson’s correlation analysis and statistical significance is denoted using * = p <0. 05, *** = p < 0.001 and **** = P < 0.0001. Graphs were generated using either GraphPad prism (version 10.3.1) or R studio (version 2024.04.0).

### Patient samples

Archival formalin-fixed paraffin-embedded (FFPE)-preserved tumour and background liver tissues were requested and obtained from the NHS tissue bank via the Department of Cellular Pathology at St James’s University Hospital, subject to the appropriate ethical consent at the point of surgery. Patients were specifically consented for research use of their explanted liver samples (REC reference 20/NW/0223). Anonymous details can be found in Supplementary table S1.

### DNA extraction

Sections of FFPE tissue were scraped using a size 22 scalpel (Swann-Morton). DNA was extracted using the MagMax FFPE ultra kit (Thermo Fisher), including an overnight protease digestion at 55 °C prior to inactivation at 90°C. Nucleic acids were then quantified using either the Qubit® dsDNA BR Assay Kit or the Qubit® RNA BR Assay Kit on a Qubit® 4 Fluorometer (all ThermoFisher).

### Whole exome sequencing

Genomic DNA was extracted from FFPE liver biopsies and processed using the Exome 2.0 kit (Twist Bioscience). All DNA samples were quantified using the Qubit® 4 Fluorometer before libraries were created and quality assessed on the Agilent Tapestation. Sequencing was performed on the Illumina® NovaSeq™ platform to a mean read depth of 90.8× coverage (Supplementary table S2). Sequencing data was aligned to human reference genome (build GRCh38/hg38) using bwa-mem (v0.7.17).

### Somatic variant discovery

Liver tumours were investigated for somatic variants using Mutect2. We compared tumours against matched normal liver biopsies taken from the same individual, in addition to a population germline resource to filter out common germline variants that occur in a wider population. GATK ‘LearnReadOrientationModel’ was used to filter out common orientation bias artifacts associated with FFPE samples. Only variants with ≥50 total read depth, > 20 quality score and flagged as “PASS” by GATK FilterMutectCalls were retained and used for somatic variant burden analysis.

### TMB calculation

TMB was quantified by calculating the number of non-synonymous, exonic variants per megabase of sequenced DNA (36.5 Mb). ‘NeoTMB’ was quantified by excluding variants that resulted in a START loss or STOP gain mutations from those used to calculate TMB.

### Further sequencing analysis

The CoMut plot was visualised in Google Colab python interface using CoMut (version 0.03). Gene lengths were calculated from start and end positions from ENSEMBL using GRCh37 coordinates. The The Cancer Genome Atlas Liver Hepatocellular Carcinoma dataset was used to identify previously reported common mutations (https://www.cancer.gov/tcga).

### Immunohistochemistry

FFPE tissue sections were stained with primary antibody as listed in Table 1 or antibody diluent alone as a control (Thermo Fisher) then appropriate horse-radish peroxidase (HRP)-conjugated secondary antibody (Vector Laboratories). Positive staining was visualised using ImmPACT DAB HRP Substrate kit (Vector). Slides were scanned using either an Aperio GT 450 DX scanner (Leica) or AxioScan Z.1 Slidescanner (Zeiss). For Haematoxylin and Eosin (H&E) staining, the procedure was the same as above, omitting antibody incubation and DAB visualisation, staining first for haematoxylin, then eosin (both Sigma).

**Table 1.**
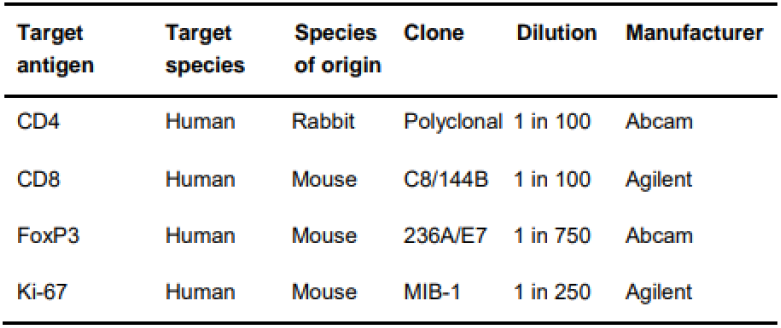
Primary antibodies used for IHC staining.

### Image analysis

Virtual slides were analysed using QuPath software version 0.5.1 (Bankhead et al., 2017). The positive cell detection feature was used to quantify the percentage of cells positive for Ki-67 staining and the number per millimetre squared of tissue for CD4, CD8 and FoxP3. A small area of each image was used for quality control and threshold parameters were adjusted between images if needed. Where possible, analysis was performed using information from the Leeds Teaching Hospital Trust’s Patient Pathway Manager Plus system to include NHS histopathology reports.

Regional analysis of Ki-67 positivity was performed on tumours where the entire section was captured on the FFPE block. Three concentric circles were created on virtual slides based on the radius of the tumour and the percentage of cells positive for Ki-67 in each circle was measured using QuPath before equations were used to calculate the Ki-67 positivity for each region (Supplementary Figure S1).

## Results

### Hepatocellular carcinoma tumours exhibit wide genetic variation

The burden of tumour-specific somatic variants in this cohort has a median value of 1.44 mutations per megabase (mut/Mb) of sequenced DNA, with significant heterogeneity observed between samples spanning over a 600-fold range (0.16 to 98.63 mut/Mb). Filtering these data for only mutations with the potential to cause nepepitope formation (neoTMB) generated similar results, with a median value of 1.44 mut/Mb and a range of 0.16 to 93.39 mut/Mb (Figure 1A).

**Figure 1.**
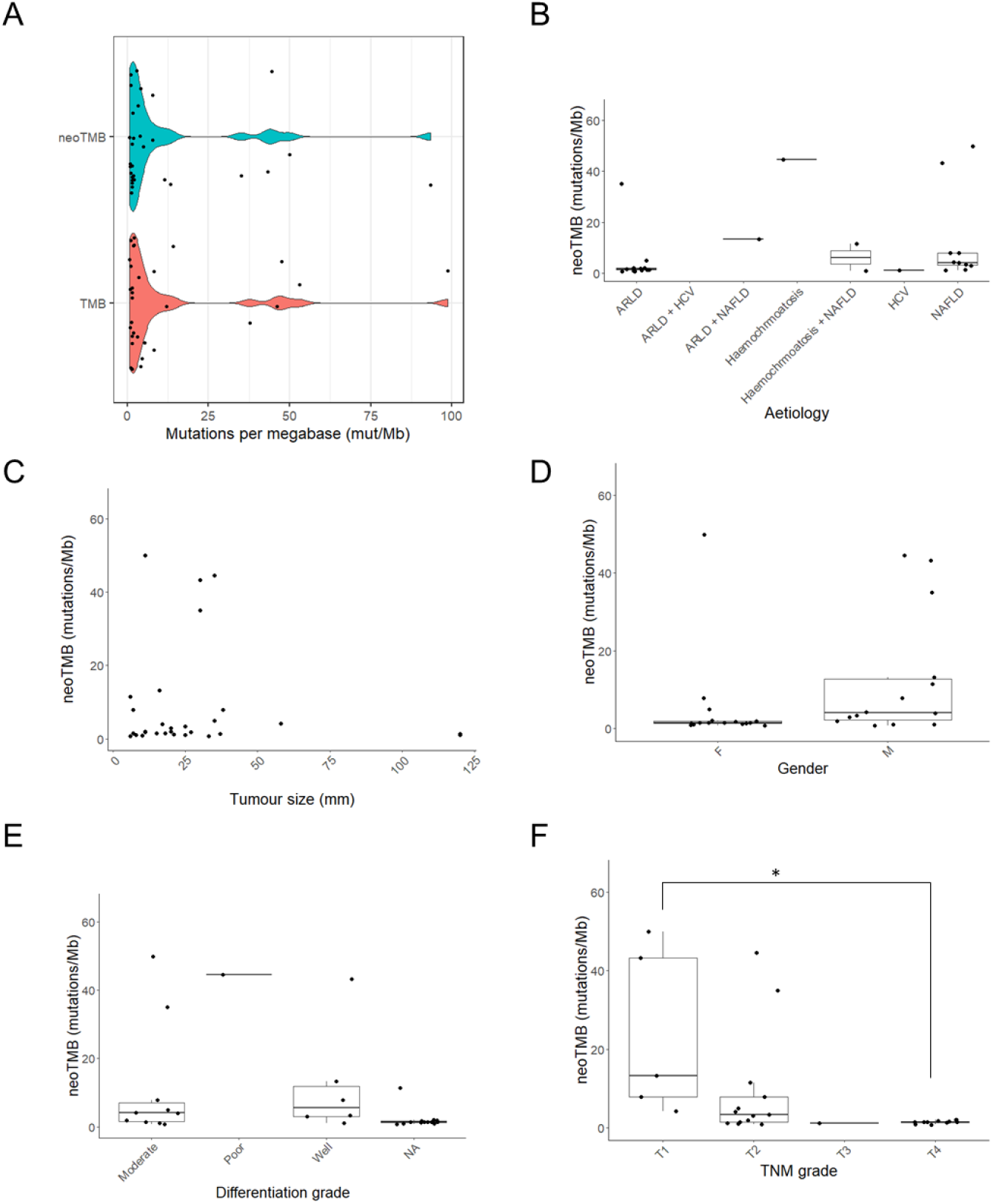
TMB and neoTMB data for hepatocellular carcinoma tumours in relation to tumour and patient characteristics. A) Distribution of TMB and neoTMB values. NeoTMB in relation to aetiological group (B), tumour size (C), gender (D), differentiation grade (E) and Tumour Node Metastasis (TNM) stage (F). Adjusted-P value represents a one-way ANOVA with Tukey correction performed (* = < 0.05). N numbers = 30 to 52, lines represent median values.

Differences in the neoTMB values were not significantly different between aetiological group (Figure 1B), tumour size (Figure 1C), gender (Figure 1D) or differentiation grade (Figure 1E). However, a significant difference was observed between the neoTMB values for stage T1 tumours when compared to stage T4 tumours, with a higher neoTMB in T1 tumours (Figure 1F). Although not reaching statistical significance, other trends that can be noted are of higher neoTMB in tumours from male patients when compared to those from female patients and also in NAFLD when compared to ARLD.

### Distinct genetic patterns found between aetiological groups

Differing mutational profiles, with respect to individual variants and specific genes containing mutations, are represented in heatmaps in Figures 2A and 2B. HCV-related tumours shared the highest degree of similarity, with some variants found in at least 50 % of tumours, and yet the same variants occurring rarely in other aetiological groups. Pathway analysis revealed no functional classification of these variants or genes, either overall or with respect to separate aetiological groups (Supplementary Tables S3-S6).

**Figure 2.**
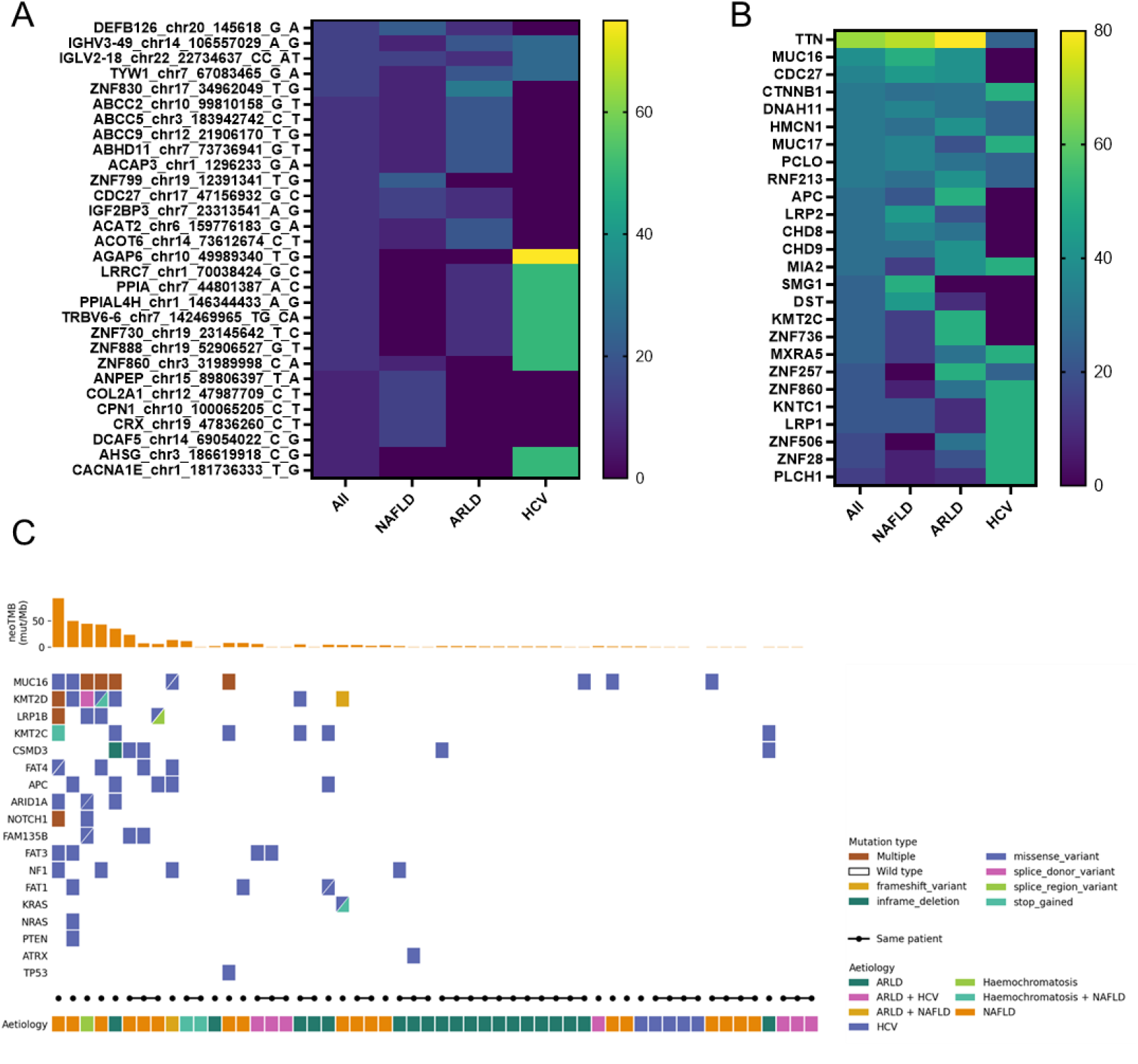
Genetic aberrations found across hepatocellular carcinoma tumours. A) Frequency of specific variants found in patients overall and within aetiological groups. B) Frequency of mutated genes found overall and within aetiological groups C) Occurrence of mutations in hepatocellular carcinoma-associated genes, according to literature reports. Multifocal tumours are indicated by joined circles. For aetiology heatmaps, mutations in multifocal tumours were only counted once per patient. N number = 4 to 28.

When looking at the cohort of tumours as a whole, there is stark variation in the genes which are mutated. Figure 2C displays a co-mut plot, where individual tumours are represented in rows, showing the most commonly mutated genes according to TCGA project cohorts of liver cancer tumours. Logically, tumours with a higher neoTMB are seen to contain a higher number of these commonly-mutated genes. Tumours from the same aetiological group or same patient do not have a noticeably higher degree of similarity when examining these genes.

Table 2A shows the most frequently mutated genes for all tumours and at an inter-patient level, counting each patient with multifocal tumours once only. The change in frequencies between these measures shows that a high proportion of multifocal tumours contain mutations in the same genes.

**Table 2.** The top 10 most commonly mutated genes in this hepatocellular carcinoma cohort. A: Mutation frequency for all tumours and on a per-patient basis. For multifocal patients, mutated genes were just counted once. B: Frequency adjusted for gene length.

| Gene | Frequency in all tumours | % frequency in all tumours | Frequency for individual patients | % frequency for individual patients |
| --- | --- | --- | --- | --- |
| TTN | 19 | 35.19 | 17 | 65.38 |
| ZNF708 | 16 | 29.63 | 6 | 23.08 |
| HMCN1 | 13 | 24.07 | 8 | 30.77 |
| CDC27 | 12 | 22.22 | 9 | 34.62 |
| PCLO | 12 | 22.22 | 8 | 30.77 |
| ZNF726 | 12 | 22.22 | 7 | 26.92 |
| DEFB126 | 11 | 20.37 | 4 | 15.38 |
| ZNF430 | 11 | 20.37 | 3 | 11.54 |
| ZNF595 | 11 | 20.37 | 5 | 19.23 |
| MUC16 | 10 | 18.52 | 10 | 38.46 |

*Table 2: The top 10 most commonly mutated genes in this hepatocellular carcinoma cohort.*
| Gene | % frequency in all tumours | Gene size (Mb) | Relative frequency (frequency/Mb) |
| --- | --- | --- | --- |
| IGLV2-18 | 15.38 | 0.0005 | 8299 |
| TRBV6-5 | 15.38 | 0.0005 | 7984 |
| PPIAL4H | 11.54 | 0.0008 | 3953 |
| HERC2 | 26.92 | 0.0020 | 3448 |
| MT-CYB | 11.54 | 0.0011 | 2632 |
| USP17L1 | 11.54 | 0.0016 | 1884 |
| PRKACG | 11.54 | 0.0016 | 1865 |
| RBMXL3 | 19.23 | 0.0035 | 1446 |
| OLIG3 | 11.54 | 0.0022 | 1367 |
| ZNF830 | 11.54 | 0.0022 | 1341 |

To account for the possibility that gene length might influence mutation frequency, the gene size was used to normalise the inter-patient frequency. The genes with the highest relative frequency are shown in Table 2B. Genes at the top of this include *IGLV2-18* and *TRBV6-5*, which encode the variable regions of IgG light chains and T cell receptors, respectively. An ‘inflation’ can be seen, where short genes are over-represented.

### Hepatocellular carcinomas grow more rapidly at the tumour periphery

Quantification of cellular proliferation confirmed that tumours have significant higher proportions of dividing cells, when compared to paired liver tissue (Figure 3Ai). When the entire tumour was captured on the pathology block, analysis of concentric regions (Supplementary Figure S1) revealed a significant increase in the percentage of dividing cells in the tumour periphery when compared to both core and intermediate regions of the same tumour (Figure 3Aii). Example IHC images can be seen in Figure S3.

**Figure 3.**
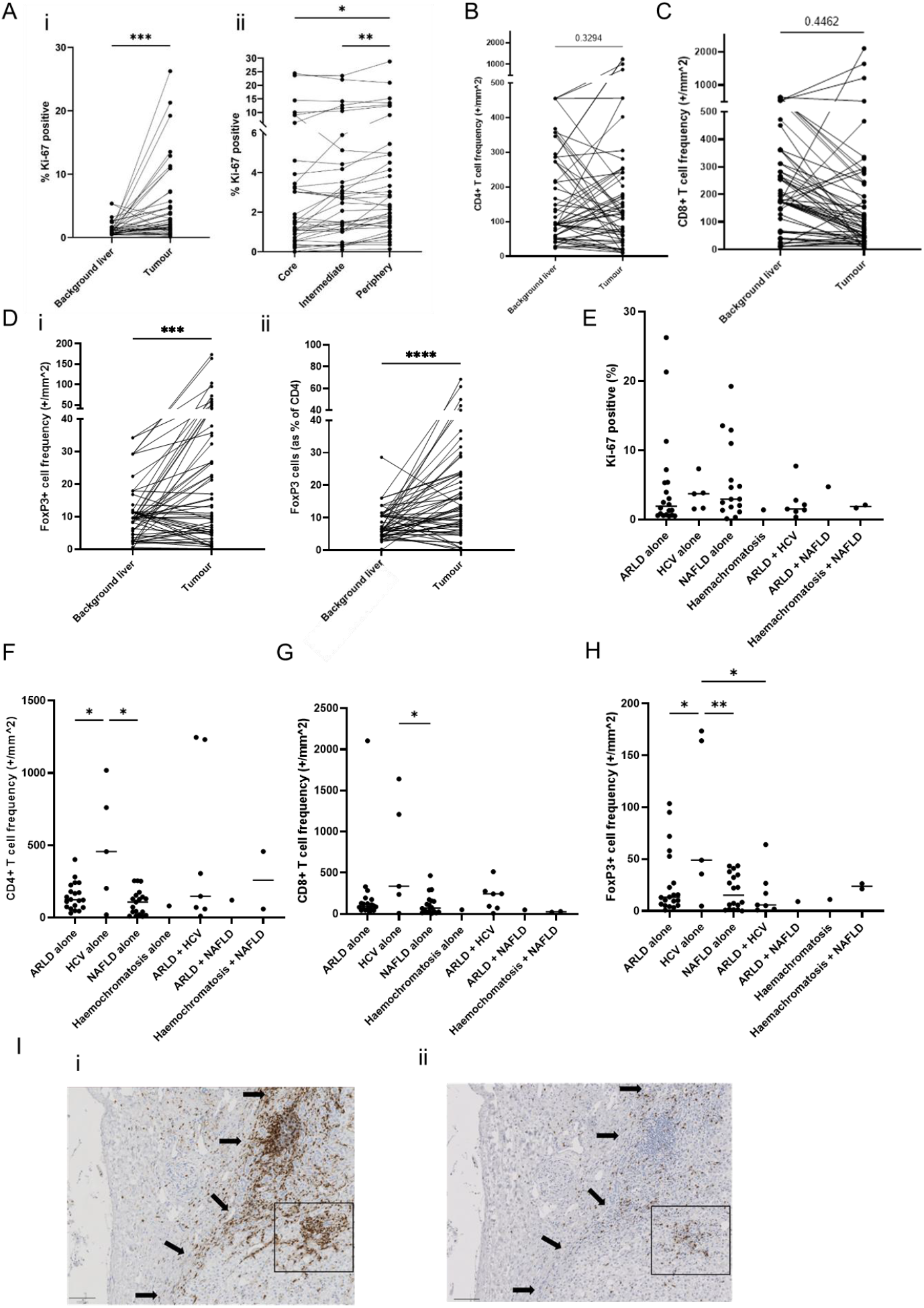
Ki-67 positivity and T cell frequency in paired background and HCC samples. A) Ki-67 positivity as a percentage of total cells in the paired liver and hepatocellular carcinoma samples (i) and between concentric regions of the same tumour (ii). Frequency of CD4+ T cell (B), CD8+ T cell (C) and FoxP3+ T cells (Di) in paired background liver and hepatocellular carcinoma tissue samples. Dii) The proportion of FoxP3+ cells as a proportion of total CD4+ T cells in paired background liver and hepatocellular carcinoma tissue samples. E) Ki-67 positivity, F) CD4+, G) CD8+ and FoxP3+ T cell frequencies in tumours separated by aetiological group. I) Representative images of CD8+ (i) and CD4+ (ii) infiltration in a HCV-related tumour. P values represent paired t-tests (* = p < 0.05, ** = p < 0.01, *** = p < 0.001 and **** = P < 0.0001) in sections A to C and One-way ANOVA in sections E to H. N numbers = 30 to 52

### Enrichment of FoxP3 but not of CD4+ or CD8+ T cells in tumours

There was no significant increase in the frequency of CD4+ or CD8+ T cells in tumours, when compared to section of background liver from the same patient. There is a large range in the number of tumour-resident cells between tumours, representing a greater than 200-fold difference in both instances (Figure 3B and C).

There was, however, a significant increase in the number of FoxP3+ T cells in tumours in comparison to the paried background liver, seen in Figure 3Di. To account for these cells also being CD4+, the frequency was also plotted as a proportion of CD4+ cells. A greater level of significance was observed in Figure 3Dii, indicating this increase is independent of the CD4+ frequency in tumours.

No correlation was found between immune cell frequencies and Ki-67 or neoTMB values (Supplementary Figure S2).

### Hepatitis C virus-associated tumours have higher T cell infiltration

The Ki-67 quantification indicated that proliferation rates were seen to be similar between aetiological groups, as seen in Figure 3E.

Patients with HCV-associated disease had a higher number of tumour-resident CD4+ (Figure 3F), CD8+ (Figure 3G) and FoxP3+ (Figure 3H) T cells when compared to tumours of other aetiological subtypes; this reached significance when compared to the NAFLD group for each cell type and also for ARLD in the case of CD4+ and FoxP3+ T cells (Figures 3F and 3H). This trend is supported by observing the non-significant increase in median value for CD4+ and FoxP3+ cells for the combined ARLD + HCV group (Figures 3F and 3H). Histology slides revealed aggregation of T cells in HCV-related patient tumours, as seen for CD4+ and CD8+ in Figure 3Ii and ii.

### Variation in tumour proliferation rate and frequency of immune cells observed between segments of the liver

Figure 4Ai shows the distribution of tumour occurrence by liver segment for those cases where the liver histology was stated on the pathology report. Segments four and seven were the most frequent locations, with no tumours located in segment one. Overall, more tumours were found in the right lobe (88.7%), when compared to the left lobe (11.3%).

**Figure 4.**
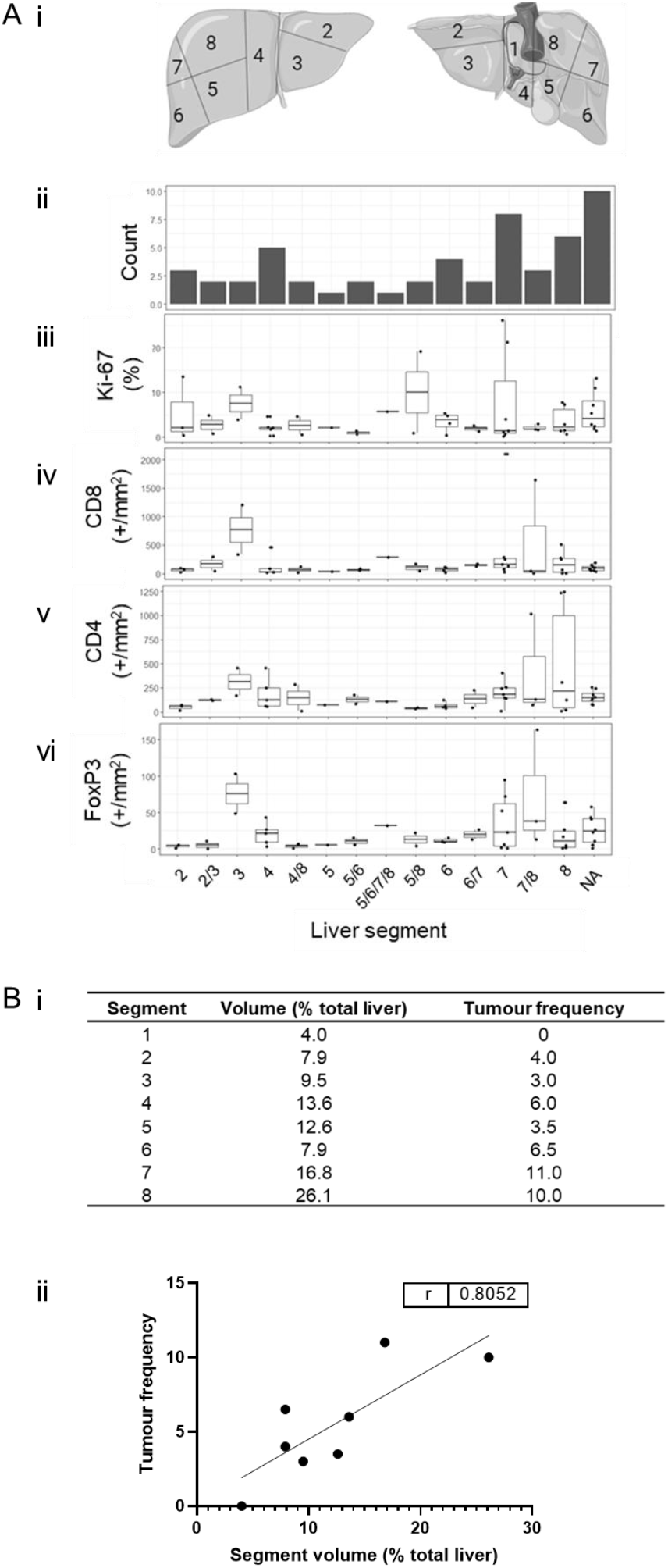
Proliferation, T cell frequency and tumour occurrence according to segments of the liver. A) Diagram of liver segments (i), segmental location of HCC tumours (ii), Ki-67 positivity (iii), frequency of tumour-infiltrating CD8+ (iv), CD8+ (v) and FoxP3+ T cells (vi) by segmental location. Liver segment information was identified from pathology reports by Leeds Teaching Hospitals Trust histopathologists, where available. NA: tumour location not available. N = 51. B) Volume of liver segments according to Mise et al. (2014) alongside tumour occurrence in each segment (i). Correlation analysis between liver segment volume and tumour frequency (ii). R value indicates the result of a Pearson’s correlation analysis.

Although this sample size does not allow for statistical evaluation, data in Figure 4Aii-vi shows that tumour proliferation and immune cell frequencies vary according to the segment carrying the tumour. Segments three, seven and eight appear to have a trend towards a higher frequency of immune cells when compared to the other segments.

Utilising data from a publication by Mise et al. in 2014 where the average volume of liver segments was modelled from CT scans of 107 normal livers (Figure 4Bi), we found a strong correlation between tumour segment location and the average volume of each segment, as seen in Figure 4Bii.

### Multifocal tumours

Of the 10 patients with multifocal tumours, most cases were spread throughout segments of the liver and, in some instances, across the two main lobes. Maximum parsimony analysis to generate phylogenetic trees indicated a range of genetic relatedness contexts. Often, within the same patients, multifocal tumours showed differing levels of genetic relatedness. Figure 5A shows one patient with four tumours across both main lobes of the liver (i), with three of these having 2 variants in common whilst tumour (T)3 is completely unrelated (ii). T2 and T4 are most genetically similar and yet have varied frequencies of FoxP3+ and CD4+ cells (iii) and are remarkably different in size (iv). T1 and T4 are most similar in immune cell frequencies, except for CD8 T cells, and are located closely within the liver.

**Figure 5.**
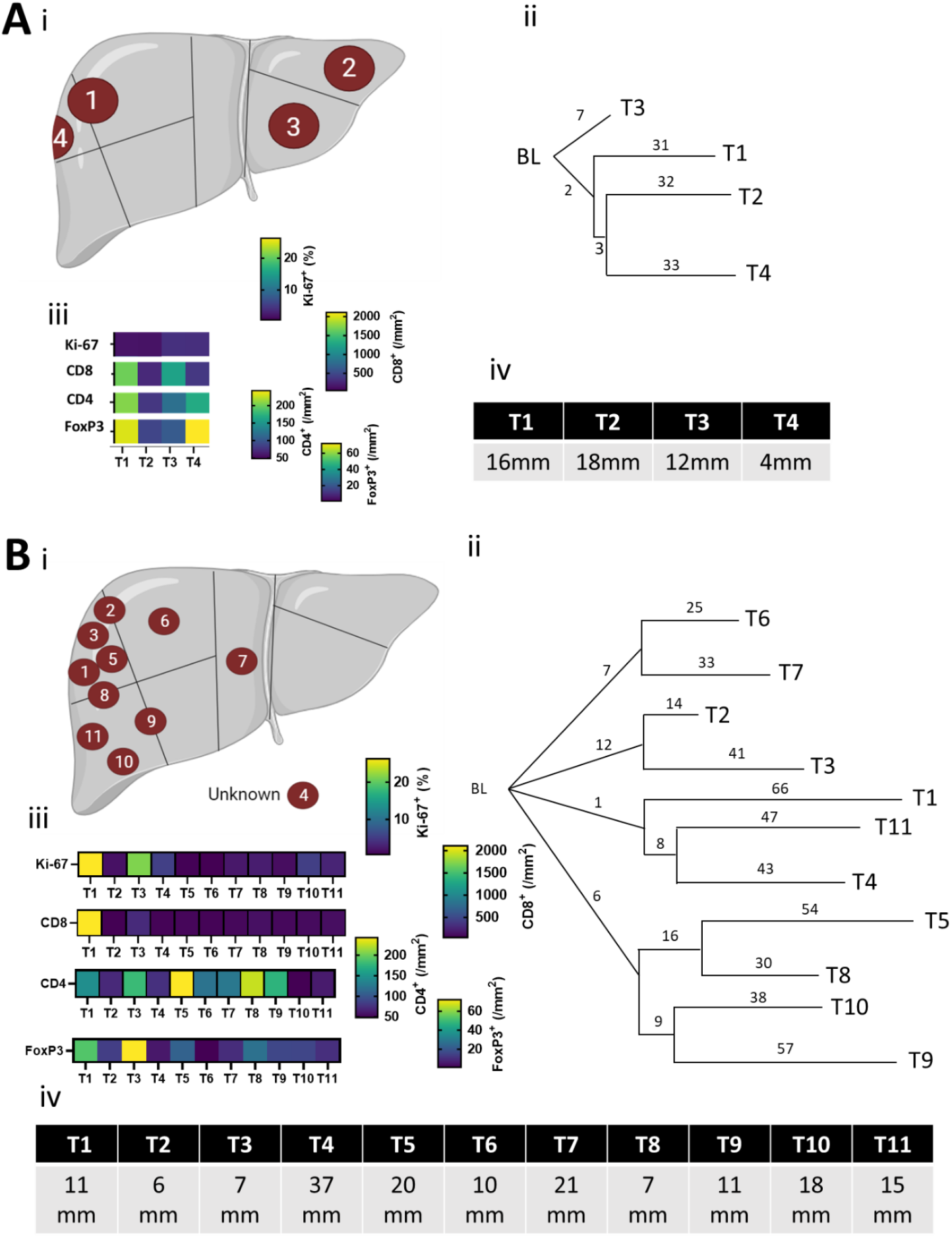
Immunogenetic relatedness of multifocal hepatocellular carcinoma tumours. Data for two patients (A and B) to indicate the segmental location of tumours (i), genetic similarity according to a Camin-Sokal phylogenetic tree analysis (ii), tumour proliferation and immune cell frequencies (iii) and tumour sizes (iv).

A patient with 11 tumours was examined as part of this analysis (Figure 6Bi). The phylogenetic tree shows that these tumours fall into 4 clades and then each tumour contains between 14 and 66 unique variants. Some clades are immunologically similar, such as T6 and T7 which are in neighbouring segments of the liver. Other clades exhibit wide variation in proliferation and immune cell frequency, such as T2 and T3. Tumours in the same clade were found in neighbouring segments for this patient. Diagrams of the other 8 patients with multifocal tumours can be seen in Supplementary Figure S4.

**Figure 6.**
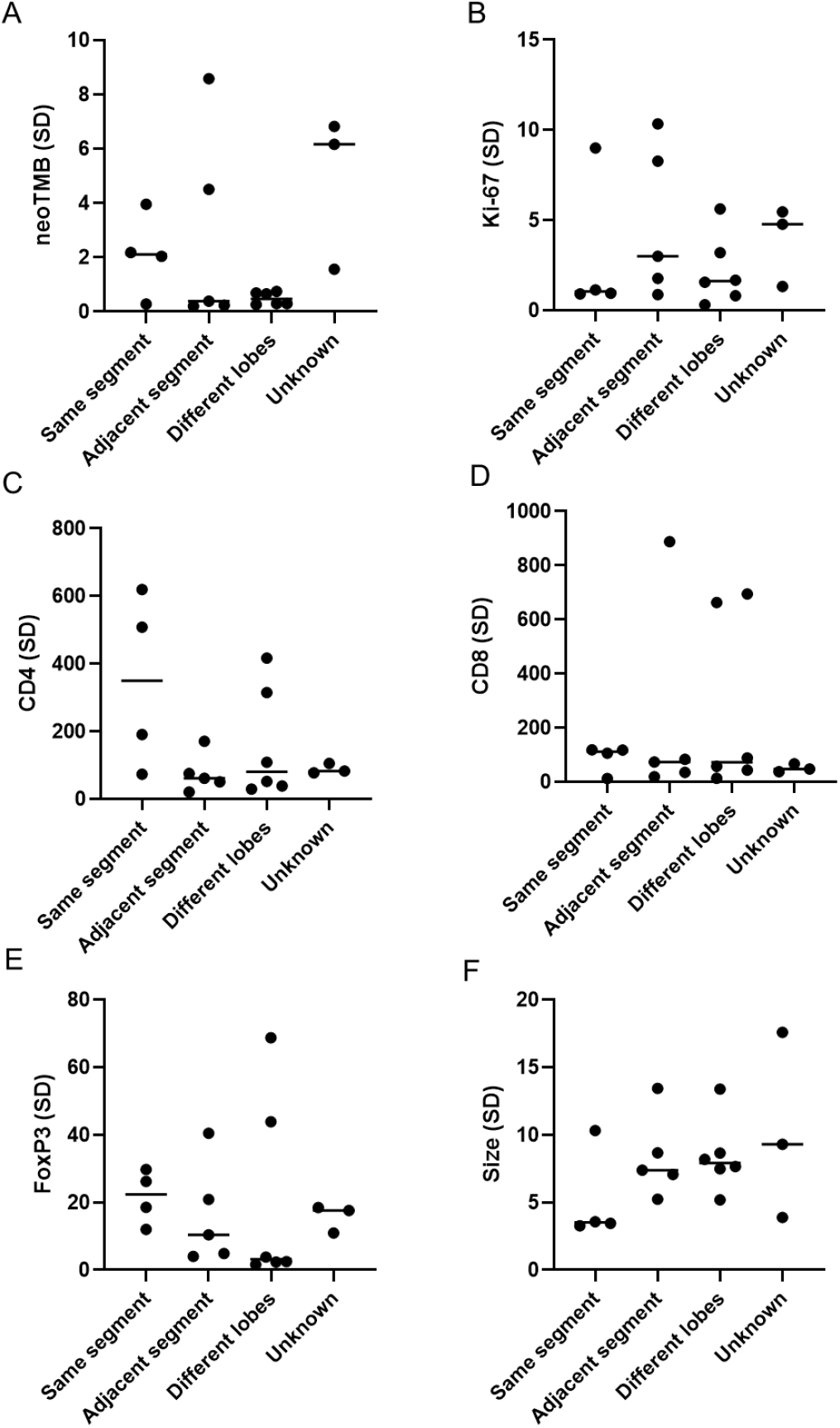
Standard deviation in tumour proliferation and immune cell frequencies in multifocal hepatocellular carcinoma tumours, according to segmental relationship. A) neoTMB, B) Ki-67 positivity, C) CD4+ T cells, D) CD8+ T cells, E) FoxP3+ T cells and F) Tumour size. Distance is categorsied by two tumours being recorded in the pathology report as being in the same segment, immediately neighbouring segments (adjacent), in the opposite lobe (different lobe) or marked as unknown if the segmental location was not recorded.

Next, the similarity of tumours depending on their spatial closeness was evaluated in Figure 6. There was no obvious trend for neoTMB (Figure 6A), Ki-67 (Figure 6B), CD4+ T cell (Figure 6C), CD8+ T cell (Figure 6D), FoxP3+ T cell (Figure 6E) or tumour size (Figure 6F). This indicates that the closeness of tumours does not make them more likely to be similar in any of these aspects.

## Discussion

This immunogenetic study of HCC tumors represents the main aetiological groups within the UK population, and demonstrates the high level of heterogeneity found within . The sequencing data generated from these tumours revealed a median TMB and neoTMB values of 1.44 mut/Mb, shown in Figure 1, which is lower than previously reported values in the literature (3.6 mut/Mb). However, our data has also highlighted that conderable range in mutation frequency can occur in tumour both between and within HCC cases. In that respect, our TMB estimations fall within the range of other reports.

The mutational profiles observed between aetiological groups in Figure 2 allude to different mutagenesis processes arising from exposure to distinct risk-factors. Whilst no conclusions could be made from the pathway analysis, the potential causative nature of these mutations warrants further investigation. For instance, *SMG1* was exclusively found to be mutated in NAFLD-associated tumours. There is no evident link between obesity and SMG1 deregulation; however, published literature does associate SMG1 haploinsufficiency with tumorigenesis in multiple organs, including in the liver (Roberts et al., 2012). The overview of commonly mutated genes within this dataset, illustrated by the co-mut plot in Figure 2C, is concordant with previous research. Observing a high frequency of mutations in the short genes for *IGLV2-18* and *TRBV6-5*, which encode the variable regions of IgG light chains and T cell receptors, could point to the concept whereby genes with critical functions are frequently mutated as this is beneficial for tumour evolution; if this happens to be a small gene then it will appear as having a high frequency.

The immunohistochemitstry data presented in Figure 3 evidences the well-recognised fact that tumours are highly proliferative and supports a model of boundary-driven growth in HCC. Our data shows that HCC tumour cells at the periphery have accelerated proliferation, a behaviour also observed in other cancer types (Bastola et al., 2020). Recently published RNA sequencing data supports this observation, but this is the first reported evidence occurring at the proteomic level (Lewinsohn et al., 2023). This growth pattern contrasts with examples of selective growth and irregular proliferation at the tumour edge which occur in colorectal cancer (Jass et al., 1996). One hypothesis is that tumour cells at the periphery have superior access to oxygen and nutrients from the tumour microenvironment (TME), to fuel higher replication rates. This concept is supported by the observation of necrotic cores in some tumours from this dataset. Higher cell replication rates, in turn, accelerate mutation rate at the periphery, which has been observed in renal cell carcinoma (Hoefflin et al., 2016).

Comparison of tumours at the aetiological level revealed that patients with hepatitis-associated tumours have a significantly higher number of tumour-resident CD4+ and CD8+ T cells, as per Figure 3. This analysis was limited to five tumours found in two patients, so does need to be interpreted with caustion; however, the observed higher frequency in tumorus from the ARLD + HCV group compared to ARLD alone adds strength to this observation. Investigating this trend in a larger group of patients is needed to validate this theory. This data may support findings from some trials that indicate patients with viral aetiologies have a better response to immunotherapy (Haber et al., 2021), although a meta-analysis shows the opposite when considering data from trials of all immune checkpoint inhibitors in HCC to date (Meyer et al., 2023). This discrepancy between T cell infiltration and response to immunotherapy raises questions as to whether there are other factors limiting the efficacy of the anti-tumour response in patients with hepatitis-associated HCC. Published work has demonstrated that the presence of an HCV viral replicon alters the repertoire of peptides presented on the surface of an HCC cell line, through modulation of the immunoproteosome (Oh et al., 2016) which could be, in part, responsible for this apparent resistance.

The number of CD4+ or CD8+ T cells in tumours were not higher than those in the background liver from the same patients. This is important when considering that the mechanism of action for current immunotherapy for advances HCC works to increase the efficacy of an existing immune response.

The concept of intrinsic resistance, whereby the therapeutic effect of a treatment is limited before the patient starts treatment (Kelderman et al., 2014), could explain why a substantial proportion of patients do not experience tumour shrinkage following atezolizumab-bevacizumab treatment.

FoxP3+ regulatory T cells have previously been shown to be significantly associated with reduced OS, in a Chinese HCC cohort (Tu et al., 2016). The data presented here supports this observation, in a different patient population, demonstrating a significant increase in the frequency of FoxP3+ T cells in the HCC tumours compared to the background liver and the further observation that this is independent of CD4+ T cell frequency. This data evidences the suppressive nature of the HCC tumour immune microenvironment and this cell subset likely limits anti-tumour immune responses and potentially confers resistance to immunotherapy treatments.

One unexpected and interesting observation is that the frequency of immune cells appears to be influenced by the segment that the tumour is in. A clear trend in Figure 4 is that tumours in segments three, seven and eight have a higher frequency of T cells. Of these, segment three is located in the left lobe whilst segments seven and eight are in the right lobe. The finding of fewer tumours being located in the left lobe is consistent with both this being the smaller of the two main lobes and with respect to previous literature. A report from a cirrhotic population found that 72.7% of HCC tumours were found in the right lobe (Renzulli et al., 2022). The same study also found that segment one was the least frequent location, which also fits with the data presented herein. The strong correlation between segmental volume and tumour occurrence supports a volume-related model of HCC distribution in the liver. Segments of the liver receive differential blood supplies from both branches of the portal vein and hepatic artery. Actual segmental blood supply varies widely between patients due to unpredictable branching of vessels (Mise et al., 2014). Despite this, to date, there is no published literature on differences in immune cell frequencies between segments. It is reasonable to consider that the vascular accessibility of the tumour to immune cells, dictated by blood flow, plus access to signalling factors, could underlie differences suggested in these data. A study to specifically look at this, with appropriate statistical power, would reveal if this pattern is a true feature of the disease.

With regards to statistical limitations in this study, it is important to note that our cohort has a male bias with a 4.7-greater frequency of male patients than females, which is higher than the population-wide frequency (Bray et al., 2018). Samples were selected based on the most recently acquired FFPE-preserved blocks, to minimise temporal nucleic acid degradation (Bass et al., 2014), and were not controlled to statistically represent a particular HCC patient population. Surgical eligibility criteria will have also influenced the demographics of the patients within this dataset; this is determined by the number and size of suspected HCC tumours, hence a skew towards a lower tumour stage is observed. Multifocal tumours are an exception and in some case histopathology forms indicate more tumours were found than suspected from MRI scans. In this set, 35.71% of patients had multifocal disease, which is likely to be an underestimate compared to the population rate due to the confounding effect of surgical eligibility criteria.

As the samples were collected in an opportunistic manor, as described above, there was no controlling for patient characteristics such as aetiology or tumour node metastasis (TNM) grade. In turn, this means that statistical analyses between groups were not always possible due to insufficient numerical power.

Multiple tumours from the same liver are varied in the frequency of immune cells and the rate at which they grow. This is seen in patients where tumours are demonstrated to share mutations that indicate metastatic evolution, in Figure 5. Published work using RNA sequencing data to infer characteristics of the TME have also noted variation in T cell infiltration and, in other populations, between multifocal HCC tumours (Xu et al., 2019). Confirming this finding at the proteomic level evidences the extent of intra-patient heterogeneity in multifocal HCC. The presence of multiple tumours with differing frequencies of immune cells is a likely mechanism of intrinsic resistance to the immunotherapy treatments currently used to treat advanced HCC. If there is variation in the immunogenicity of the tumours within the same patient, this is likely to result in a disproportionate anti-tumour response, with some tumours shrinking but others able to continue proliferating, to limit overall treatment efficacy.

Taken together, the low mutation-driven immunogenicity, poor immune cell infiltration and heterogeneity in multifocal disease, this data supports the need for therapeutic approaches that broaden the adaptive anti-tumour immune response. These could include personalised vaccination strategies, oncolytic viruses and strategies for modulating the immunosuppressive TME; all of which could be used in combination with existing standard of care immune checkpoint blockade. Future work in larger, aetiology-stratified cohorts with integrated genomic and spatial immune profiling will be essential to validate these observations.

## Acknowledgments

We are grateful to all the patients that provided consent for use of samples in research. The research is supported by the National Institute for Health Research (NIHR) infrastructure at Leeds. The views expressed are those of the author(s) and not necessarily those of the NHS, the NIHR or the Department of Health.

## Data availability

Further enquiries can be directed to the corresponding author.

## List of abbreviations

ARLD: Alcohol-related liver disease
DAB: Diaminobenzidine
DNA: Deoxyribonucleic acid
FFPE: Formalin-fixed paraffin embedded
GATK: Genome analysis toolkit
HBV: Hepatitis B virus
HCC: Hepatocellular carcinoma
HCV: Hepatitis C virus
HRP: Horseradish peroxidase
ICB: Immune checkpoint blockade
MRC: Medical Research Council
MRI: Magnetic resonance imaging
NAFLD: Non-alcoholic fatty liver disease
NHS: National Health Service
NIHR: National Institute for Health and Care Research
REC: Research Ethics Committee
RNA: Ribonucleic acid
TCGA-LIHC: The cancer genoma atlas liver hepatocellular carcinoma
TMB: Tumour mutation burden
TME: Tumour microenvironment
TNM: Tumour Node Metastasis
TTN: Titin

