## Supplementary data for "The immune and genetic heterogeneity of hepatocellular carcinoma, with a focus on multifocal disease"

**Supplementary Table 1**: *Characteristics of patient samples, as per the NHS pathology reports.*

| Patient ID | Gender | Aetiology | TNM grade | No. of tumours |
|---|---|---|---|---|
| 13 | M | NAFLD | T2 | 1 |
| 14 | M | NAFLD | T2 | 1 |
| 25 | M | ARLD | T3 | 3 |
| 44 | M | NAFLD | T1 | 1 |
| 52 | M | HCV | T4 | 4 |
| 725 | M | ARLD | T2 | 2 |
| 728 | M | ARLD + HCV | T2 | 3 |
| 97 | M | ARLD + NAFLD | T1 | 1 |
| 10 | F | NAFLD | T3 | 1 |
| 17 | M | ARLD | T2 | 1 |
| 220 | M | ARLD | T2 | 1 |
| 221 | M | Haemochromatosis | T2 | 1 |
| 222 | M | NAFLD | T2 | 2 |
| 223 | M | Haemochromatosis + NAFLD | T2 | 2 |
| 224 | F | ARLD | T4 | 11 |
| 225 | M | HCV | T2 | 1 |
| 226 | F | NAFLD | T4 | 4 |
| 227 | M | NAFLD | T2 | 1 |
| 228 | M | ARLD + HCV | T4 | 4 |
| 229 | F | ARLD | T2 | 1 |
| 230 | F | NAFLD | T1 | 1 |
| 231 | F | NAFLD | T2 | 1 |
| 232 | M | NAFLD | T3 | 3 |
| 233 | M | ARLD | T2 | 1 |
| 234 | M | NAFLD | T1 | 1 |
| 235 | M | UNKNOWN | T1 | 1 |
| 236 | M | NAFLD | T4 | 1 |
| 237 | M | NAFLD | T1 | 1 |

**Supplementary Table 2**: Whole exome sequencing metrics

| Sample | total | mean | granular_third_quartile | granular_median | granular_first_quartile | %_bases_above_15 |
|---|---|---|---|---|---|---|
| 10_1D_S26_L003 | 1445054523 | 38.61 | 45 | 28 | 17 | 77.7 |
| 10D_S25_L003 | 1760848602 | 47.04 | 54 | 35 | 22 | 85.9 |
| 13_1D_S2_L003 | 1936934357 | 51.75 | 68 | 45 | 28 | 90.4 |
| 13D_S1_L003 | 2769390306 | 73.99 | 89 | 71 | 54 | 97.8 |
| 14_1D_S4_L003 | 2701647779 | 72.18 | 87 | 65 | 48 | 97.5 |
| 14D_S3_L003 | 3287963619 | 87.85 | 105 | 81 | 61 | 98 |
| 17_1D_S28_L003 | 927473865 | 24.78 | 26 | 17 | 10 | 54.2 |
| 17D_S27_L003 | 2734941839 | 73.07 | 86 | 54 | 35 | 93.9 |
| 220_1_L011 | 2984663192 | 79.74 | 96 | 65 | 43 | 96.4 |
| 220_L011 | 2907114731 | 77.67 | 90 | 61 | 41 | 96.6 |
| 221_1_L011 | 2376001604 | 63.48 | 77 | 52 | 35 | 94.5 |
| 221_L011 | 1666642730 | 44.53 | 51 | 32 | 20 | 83.9 |
| 222_1_L011 | 4495057417 | 120.1 | 137 | 107 | 82 | 98.2 |
| 222_2_L011 | 3810904863 | 101.82 | 121 | 92 | 66 | 97.2 |
| 222_L011 | 5985803933 | 159.92 | 187 | 143 | 107 | 98.3 |
| 223_1_L011 | 2646635797 | 70.71 | 89 | 60 | 38 | 94.8 |
| 223_2_L011 | 5298757219 | 141.57 | 170 | 128 | 95 | 98 |

| | | | | | | |
|---|---|---|---|---|---|---|
| 223_L011 | 2008168789 | 53.65 | 63 | 46 | 32 | 95.2 |
| 224_1_L011 | 5439751787 | 145.33 | 170 | 135 | 106 | 97.9 |
| 224_10_L011 | 2773563646 | 74.1 | 93 | 50 | 19 | 78.7 |
| 224_11_L011 | 4050274187 | 108.21 | 128 | 91 | 65 | 97.6 |
| 224_2a_L011 | 2844035509 | 75.98 | 102 | 51 | 18 | 77.4 |
| 224_3a_L011 | 2772718600 | 74.08 | 100 | 57 | 25 | 83.5 |
| 224_4_L011 | 3467633587 | 92.65 | 112 | 79 | 54 | 97 |
| 224_5_L011 | 2386680991 | 63.77 | 81 | 45 | 17 | 77 |
| 224_6_L011 | 2633136537 | 70.35 | 87 | 57 | 35 | 93 |
| 224_7_L011 | 1760800754 | 47.04 | 59 | 33 | 14 | 72.2 |
| 224_8a_L011 | 2272452868 | 60.71 | 77 | 45 | 21 | 80.8 |
| 224_9_L011 | 2039145905 | 54.48 | 66 | 41 | 23 | 84 |
| 224_L011 | 4551067564 | 121.59 | 159 | 111 | 67 | 96.7 |
| 225_1_L011 | 1819292117 | 48.61 | 59 | 37 | 23 | 86.3 |
| 225_L011 | 4146928494 | 110.79 | 132 | 99 | 72 | 97.7 |
| 226_1a_L011 | 1956890128 | 52.28 | 69 | 42 | 22 | 83.5 |
| 226_2_L011 | 1971072369 | 52.66 | 65 | 43 | 27 | 90.3 |
| 226_3_L011 | 2398799056 | 64.09 | 81 | 53 | 33 | 92 |
| 226_4_L011 | 3065118273 | 81.89 | 112 | 66 | 30 | 86.6 |

| | | | | | | |
|---|---|---|---|---|---|---|
| 226_L011 | 2679485082 | 71.59 | 92 | 47 | 18 | 77.7 |
| 227_1a_L011 | 1652797394 | 44.16 | 59 | 30 | 11 | 68.3 |
| 227_L011 | 3390420061 | 90.58 | 104 | 72 | 50 | 97.6 |
| 228_1a_L011 | 2743147267 | 73.29 | 103 | 65 | 33 | 88.9 |
| 228_2_L011 | 4611512553 | 123.21 | 159 | 108 | 68 | 96.3 |
| 228_3_L011 | 3952477815 | 105.6 | 126 | 96 | 71 | 97.9 |
| 228_4_L011 | 5793099121 | 154.78 | 184 | 135 | 98 | 98.2 |
| 228_L011 | 3578885934 | 95.62 | 115 | 86 | 62 | 97.7 |
| 229_1_L011 | 2144648178 | 57.3 | 76 | 48 | 25 | 84.6 |
| 229_L011 | 5141754549 | 137.37 | 158 | 120 | 91 | 98.1 |
| 230_1_L011 | 3200015482 | 85.5 | 105 | 80 | 59 | 97.4 |
| 230_L011 | 4695952521 | 125.46 | 154 | 118 | 86 | 97.8 |
| 231_1_L011 | 3642512339 | 97.32 | 134 | 83 | 47 | 94 |
| 231_L011 | 2434296864 | 65.04 | 81 | 60 | 43 | 96.3 |
| 232_10_L011 | 3945604662 | 105.42 | 129 | 97 | 71 | 97.9 |
| 232_2_L011 | 2017501357 | 53.9 | 68 | 47 | 29 | 90.2 |
| 232_3_L011 | 2437720399 | 65.13 | 80 | 56 | 37 | 94.1 |
| 232a_L011 | 2660296760 | 71.08 | 87 | 57 | 36 | 95 |
| 233_1_L011 | 2119596444 | 56.63 | 74 | 35 | 13 | 71.4 |

| | | | | | | |
|---|---|---|---|---|---|---|
| 233a_L011 | 3755932195 | 100.35 | 120 | 92 | 68 | 97.9 |
| 234_1_L011 | 6139659161 | 164.03 | 202 | 152 | 110 | 98 |
| 234_pool_L011 | 3656356711 | 97.69 | 125 | 90 | 60 | 96.6 |
| 235_10_L011 | 5432644741 | 145.14 | 182 | 130 | 90 | 98.2 |
| 235_11_L011 | 6479946423 | 173.13 | 216 | 152 | 108 | 98.2 |
| 235_12_L011 | 5477858356 | 146.35 | 183 | 130 | 90 | 98.1 |
| 235_L011 | 1682906662 | 44.96 | 56 | 31 | 15 | 73.9 |
| 236_10_L011 | 7568776108 | 202.22 | 270 | 151 | 91 | 98.2 |
| 236_11_L011 | 3080666262 | 82.31 | 102 | 57 | 28 | 88.2 |
| 236_12_L011 | 2898900399 | 77.45 | 91 | 49 | 22 | 81.2 |
| 236_13_L011 | 3462171917 | 92.5 | 116 | 69 | 36 | 91.7 |
| 236_L011 | 2668227655 | 71.29 | 84 | 41 | 16 | 76.1 |
| 237_1_L011 | 1843156225 | 49.24 | 62 | 41 | 26 | 90 |
| 237_pool_L011 | 3290698505 | 87.92 | 115 | 73 | 43 | 94 |
| 25_1D_S6_L003 | 3442035297 | 91.96 | 109 | 86 | 65 | 98 |
| 25_2D_S7_L003 | 3656147138 | 97.68 | 110 | 83 | 62 | 98 |
| 25_3D_S8_L003 | 2217551296 | 59.25 | 70 | 54 | 40 | 97.3 |
| 25D_S5_L003 | 4745654946 | 126.79 | 151 | 118 | 88 | 98.1 |
| 44_1D_S10_L003 | 3197002842 | 85.41 | 103 | 81 | 61 | 97.5 |

| | | | | | | |
|---|---|---|---|---|---|---|
| 44D_S9_L003 | 1173527840 | 31.35 | 39 | 25 | 14 | 72.2 |
| 52_1D_S12_L003 | 5702903983 | 152.37 | 182 | 144 | 110 | 98.1 |
| 52_2D_S13_L003 | 4062377316 | 108.54 | 131 | 99 | 74 | 97.7 |
| 52_4D_S15_L003 | 4051699512 | 108.25 | 131 | 103 | 79 | 97.7 |
| 525_3D_S14_L003 | 2513208682 | 67.15 | 82 | 50 | 30 | 91.3 |
| 52D_S11_L003 | 1441983861 | 38.53 | 50 | 33 | 20 | 82.2 |
| 725_1D_S17_L003 | 1387452646 | 37.07 | 44 | 31 | 21 | 87.4 |
| 725_2D_S18_L003 | 3122194031 | 83.42 | 100 | 67 | 44 | 95.7 |
| 725D_S16_L003 | 2096247826 | 56.01 | 68 | 52 | 38 | 96.1 |
| 728_1D_S20_L003 | 1870866327 | 49.98 | 58 | 38 | 25 | 89.4 |
| 728_2D_S21_L003 | 3845098529 | 102.73 | 122 | 96 | 74 | 97.8 |
| 728_3D_S22_L003 | 1822923806 | 48.7 | 57 | 35 | 21 | 84.5 |
| 728D_S19_L003 | 1787902383 | 47.77 | 58 | 37 | 23 | 87.4 |
| 97_1D_S24_L003 | 4189599390 | 111.93 | 130 | 102 | 78 | 97.9 |
| 97D_1_S23_L003 | 3352330421 | 89.56 | 111 | 76 | 51 | 96.2 |
| J235_1_L011 | 3850764452 | 102.88 | 129 | 91 | 61 | 97.2 |
| J235_L011 | 4858422289 | 129.8 | 159 | 110 | 74 | 97.8 |
| J236_1_L011 | 4646735501 | 124.15 | 151 | 103 | 68 | 97.6 |
| J236_1b_L011 | 2088106244 | 55.79 | 69 | 46 | 30 | 92.4 |

| | | | | | | |
|---|---|---|---|---|---|---|
| J236_L011 | 3827922705 | 102.27 | 123 | 87 | 60 | 97.1 |
| J237_1_L011 | 4620870540 | 123.46 | 148 | 106 | 74 | 97.7 |
| J237_L011 | 5719555895 | 152.81 | 192 | 119 | 77 | 97.7 |
| J238_1_L011 | 11505612677 | 307.4 | 417 | 224 | 121 | 98.1 |
| J238_L011 | 5746510274 | 153.53 | 179 | 134 | 99 | 98 |
| J239_1_L011 | 4840471272 | 129.32 | 156 | 119 | 87 | 98 |
| J239_L011 | 4093555929 | 109.37 | 139 | 83 | 48 | 95.4 |

$$Core = C$$

$$Intermediate = B - C$$

$$Periphery = A - (B + C)$$

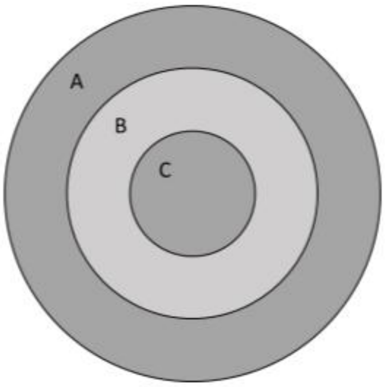


**Supplementary Figure S1:** *Diagram to show overlapping concentric regions created on virtual slides and the equations used for calculating regional Ki-67 positivity.*

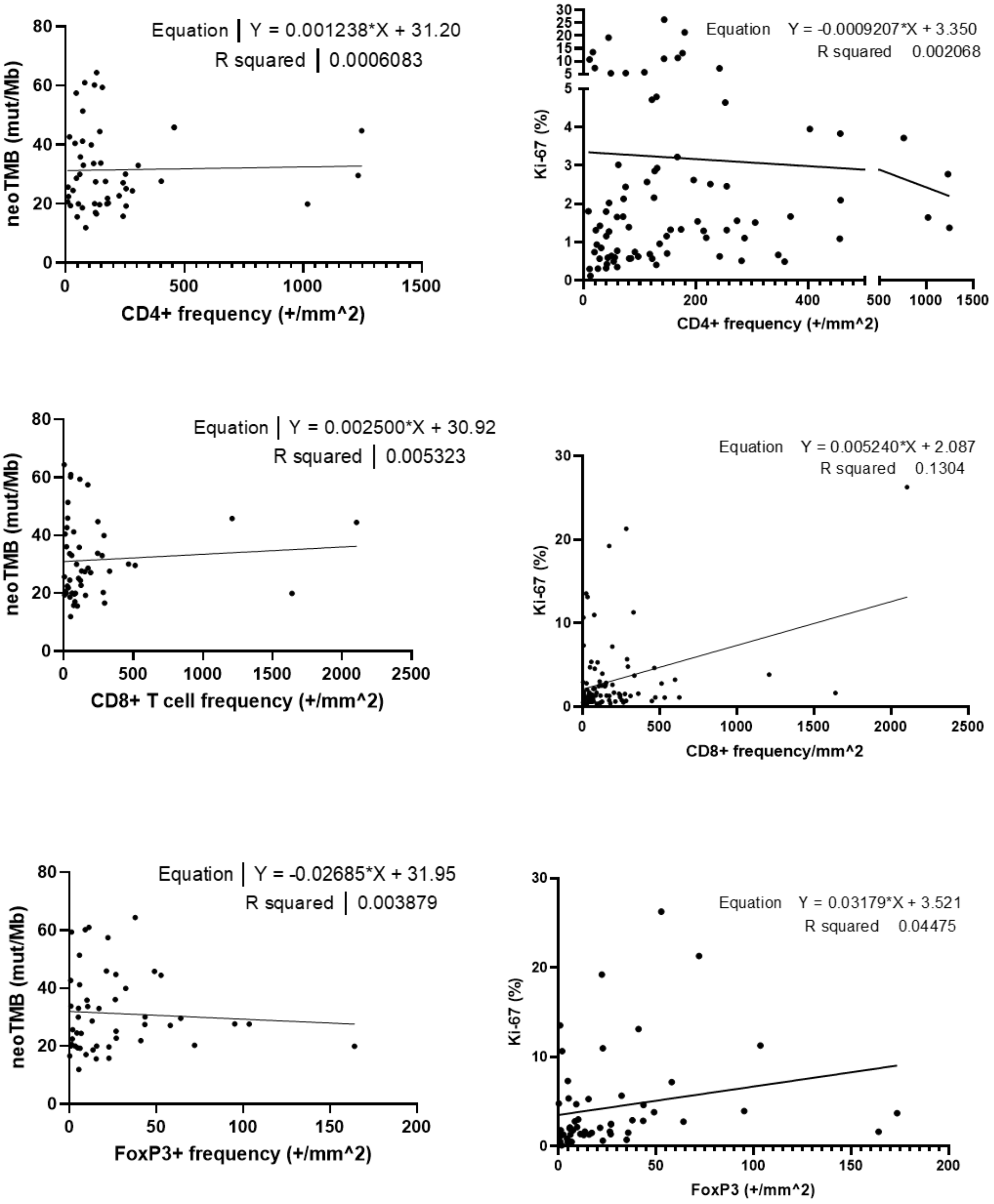


**Supplementary Figure S2:** *Relationship between neoTMB and Ki-67 or T cell frequencies*.

In each case the equation represents the line of best fit and R-squared values show the fit of the data. N = 52.

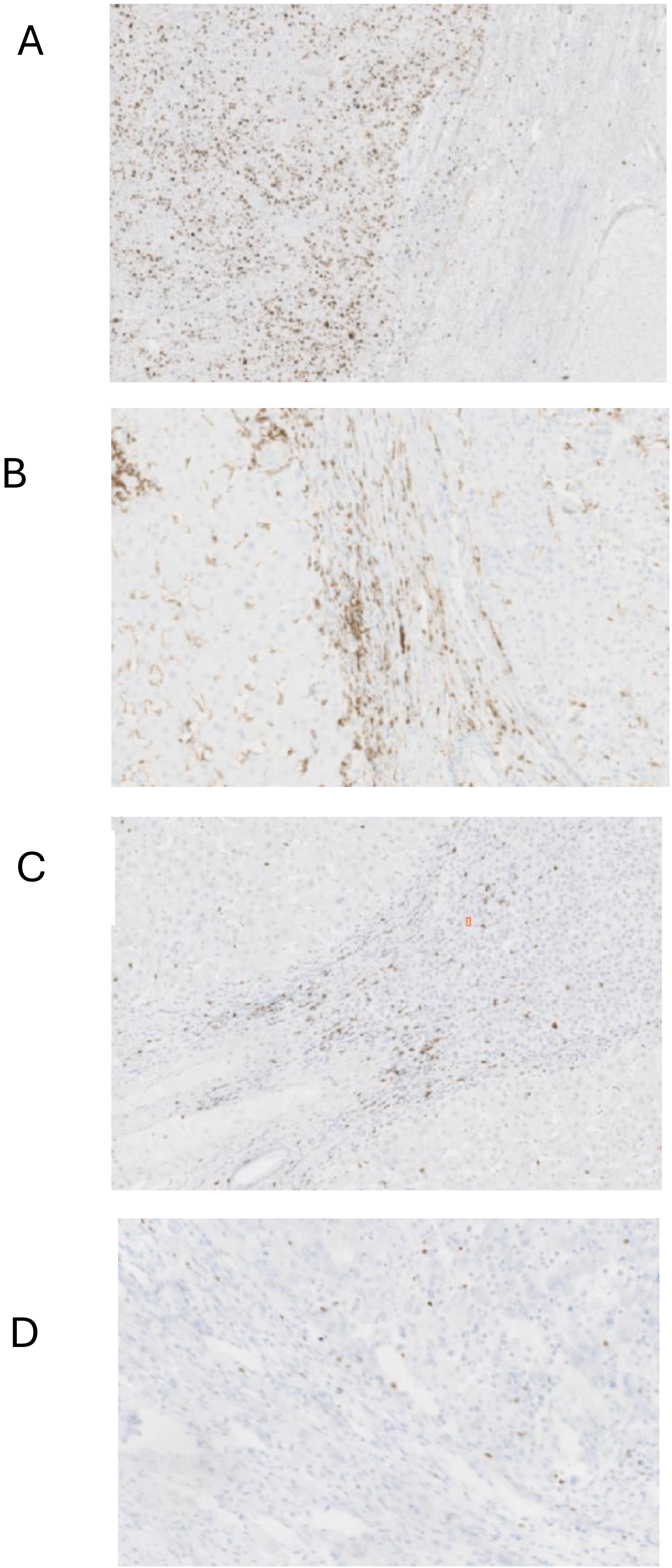


**Supplementary figure S3**: Example IHC images. A) Ki-67, B) CD4+ T cell, C) CD8+ T cell and D) FoxP3+ T cell.

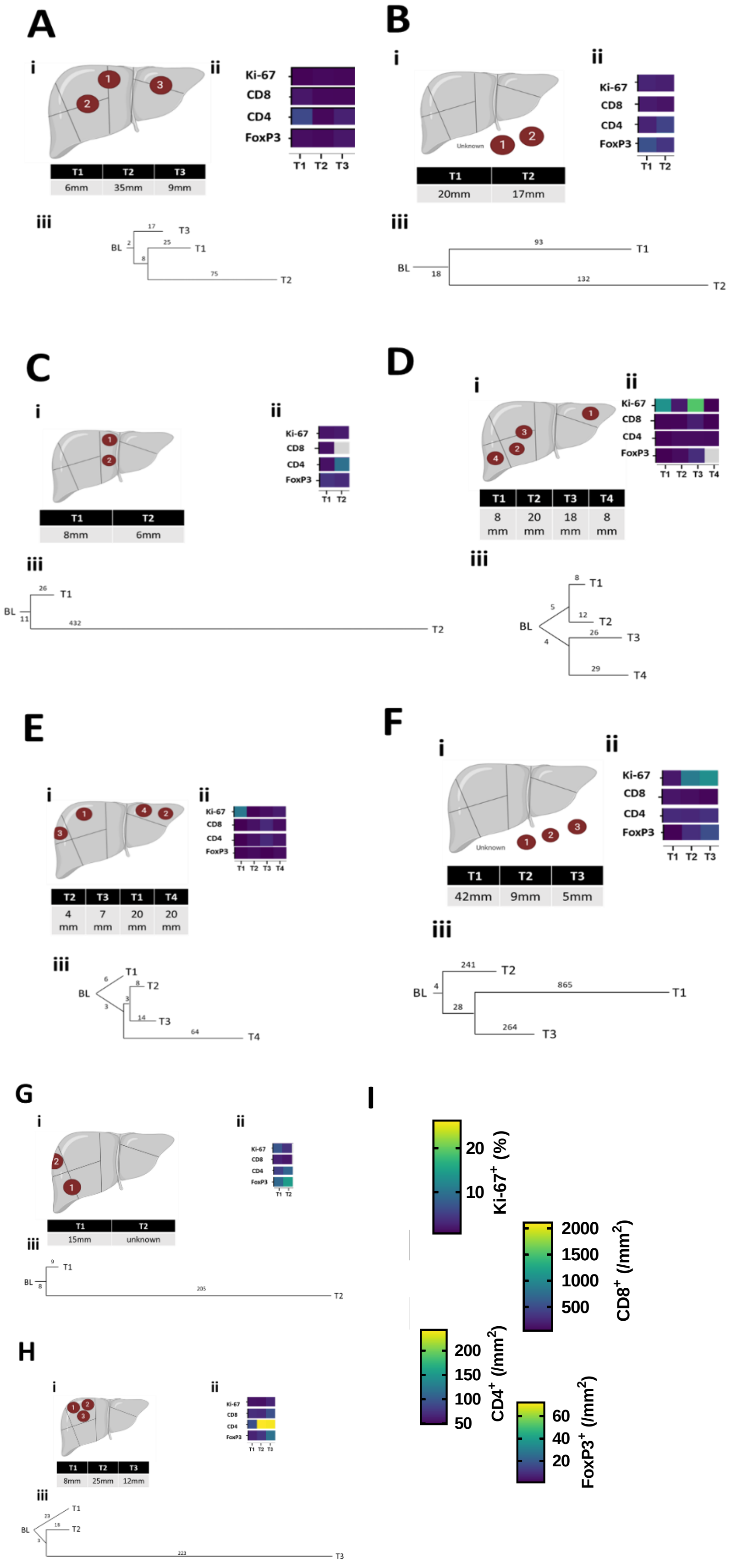

**Supplementary Figure S4:** *Characterisation of multifocal tumours*

Multifocal patients (A-H) represented with a diagram of the tumour location (i); quantification of Ki-67, CD8+, CD4+, Ki-67+ T cell immunohistochemistry (ii) and phylogenetic trees representing genetic similarity of tumours (iii). Scale bar for immunohistochemistry staining shown in I.

| source | term_name | term_id | adjusted_p_value | negative_log10_of_adjusted_p_value | term_size | query_size | intersection_size | effective_domain_size | intersections |
|---|---|---|---|---|---|---|---|---|---|
| GO:BP | detection of muscle stretch | GO:0035995 | 0.003796073 | 2.420665443 | 7 | 9 | 2 | 21026 | TTN,CTNNB1 |
| GO:BP | synaptic vesicle clustering | GO:0097091 | 0.021648547 | 1.664571247 | 16 | 9 | 2 | 21026 | CTNNB1,PCLO |
| GO:CC | presynaptic active zone cytoplasmic component | GO:0098831 | 0.002749454 | 2.560753538 | 18 | 9 | 2 | 22149 | CTNNB1,PCLO |
| GO:CC | cell cortex region | GO:0099738 | 0.01127841 | 1.947752104 | 36 | 9 | 2 | 22149 | CTNNB1,PCLO |
| GO:CC | cytoplasmic region | GO:0099568 | 0.027080454 | 1.567344062 | 313 | 9 | 3 | 22149 | CTNNB1,DNAH11,PCLO |
| GO:CC | extracellular region | GO:0005576 | 0.029610294 | 1.52855728 | 4258 | 9 | 7 | 22149 | TTN,MUC16,CTNNB1,MUC17,HMCN1,DNAH11,PCLO |
| GO:CC | cell cortex | GO:0005938 | 0.031081539 | 1.50749748 | 328 | 9 | 3 | 22149 | CTNNB1,HMCN1,PCLO |
| GO:CC | proximal portion of axoneme | GO:0120134 | 0.049918782 | 1.301736019 | 1 | 9 | 1 | 22149 | DNAH11 |
| REAC | Defective GALNT12 causes CRCS1 | REAC:R-HSA-5083636 | 0.009550725 | 2.019963656 | 16 | 7 | 2 | 11004 | MUC16,MUC17 |
| REAC | Defective GALNT3 causes HFTC | REAC:R-HSA-5083625 | 0.009550725 | 2.019963656 | 16 | 7 | 2 | 11004 | MUC16,MUC17 |
| REAC | Defective C1GALT1C1 causes TNPS | REAC:R-HSA-5083632 | 0.010820874 | 1.96573765 | 17 | 7 | 2 | 11004 | MUC16,MUC17 |
| REAC | Termination of O-glycan biosynthesis | REAC:R-HSA-977068 | 0.020093424 | 1.696946057 | 23 | 7 | 2 | 11004 | MUC16,MUC17 |
| REAC | Dectin-2 family | REAC:R-HSA-5621480 | 0.025788241 | 1.588578281 | 26 | 7 | 2 | 11004 | MUC16,MUC17 |
| CORUM | TCF4-CTNNB1 complex | CORUM:5262 | 0.049931693 | 1.301623705 | 1 | 2 | 1 | 3383 | CTNNB1 |
| CORUM | Cell-cell junction complex (CDH1-CTNNB1) | CORUM:5281 | 0.049931693 | 1.301623705 | 1 | 2 | 1 | 3383 | CTNNB1 |
| CORUM | CTNNB1-DNMT1 complex | CORUM:6336 | 0.049931693 | 1.301623705 | 1 | 2 | 1 | 3383 | CTNNB1 |

**Supplementary Table S3:** *Pathway analysis of genes mutated in at least 30% of all tumours; patients with multifocal disease are only represented once.*

| source | term_name | term_id | adjusted_p_value | negative_log10_of_adjusted_p_value | term_size | query_size | intersection_size | effective_domain_size | intersections |
|---|---|---|---|---|---|---|---|---|---|
| GO:MF | ion binding | GO:0043167 | 1.68954E-06 | 5.772230283 | 6377 | 147 | 81 | 20246 | TTN,KMT2C,ZNF736,ZNF257,HMCN1,RNF213,CHD9,KIF14,CTCF,USP32,TYW1,ITGA2,PRKDC,ZNF66,KDM6A,ZNF431,DNAH11,PCLO,CHD8,ZNF721,CACNA1H,FLT4,RYR1,ZNF726,ZNF729,DYNC2H1,RYR3,ZNF761,ZNF845,ZNF93,CACNA1A,CELSR2,CELSR3,DNAH3,DNAH5,GNL1,KIFC2,MTF2,NF1,UTRN,ZMIZ2,ZNF708,DOCK1,ZNF595,ZNF765,ZNF98,CELSR1,DGKK,EHD2,GTF2E1,IP6K2,KIF7,MST1R,PLA2G4C,PRKCG,RIN2,SLC25A23,ZFHX4,ZNF830,ZNF91,ZNF860,ZNF737,ADAMTS10,BTAF1,COLEC12,CPXM1,EGFLAM,METAP1,NLRP4,ZNF429,ZNF610,ZNF718,ZNF728,ZNF90,ZNF506,DOCK5,GFM2,IDH1,NELL2,RAB21,ZNF141 |
| GO:MF | RNA polymerase II cis-regulatory region sequence-specific DNA binding | GO:0000978 | 4.39751E-06 | 5.356793308 | 1183 | 147 | 29 | 20246 | ZNF736,ZNF257,CTCF,ZNF66,KDM6A,ZNF431,ZNF721,ZNF726,ZNF729,ZNF761,ZNF845,ZNF93,ZNF708,ZNF595,ZNF765,ZNF98,ZFHX4,ZNF91,ZNF860,ZNF737,PAX8,ZNF429,ZNF610,ZNF718,ZNF728,ZNF90,ZNF506,HSF5,ZNF141 |
| GO:MF | cis-regulatory region sequence-specific DNA binding | GO:0000987 | 6.75095E-06 | 5.170635206 | 1206 | 147 | 29 | 20246 | ZNF736,ZNF257,CTCF,ZNF66,KDM6A,ZNF431,ZNF721,ZNF726,ZNF729,ZNF761,ZNF845,ZNF93,ZNF708,ZNF595,ZNF765,ZNF98,ZFHX4,ZNF91,ZNF860,ZNF737,PAX8,ZNF429,ZNF610,ZNF718,ZNF728,ZNF90,ZNF506,HSF5,ZNF141 |
| GO:MF | small molecule binding | GO:0036094 | 8.22134E-06 | 5.08505754 | 6577 | 147 | 81 | 20246 | TTN,KMT2C,ZNF736,ZNF257,HMCN1,RNF213,CHD9,KIF14,CTCF,USP32,TYW1,ITGA2,PRKDC,ZNF66,KDM6A,ZNF431,DNAH11,PCLO,CHD8,ZNF721,CACNA1H,FLT4,RYR1,ZNF726,ZNF729,DYNC2H1,RYR3,ZNF761,ZNF845,ZNF93,CACNA1A,CELSR2,CELSR3,DNAH3,DNAH5,GNL1,KIFC2,MTF2,NF1,UTRN,ZMIZ2,ZNF708,DOCK1,ZNF595,ZNF765,ZNF98,CELSR1,DGKK,EHD2,GTF2E1,IP6K2,KIF7,MST1R,PLA2G4C,PRKCG,RIN2,SLC25A23,ZFHX4,ZNF830,ZNF91,ZNF860,ZNF737,ADAMTS10,BTAF1,COLEC12,CPXM1,EGFLAM,METAP1,NLRP4,ZNF429,ZNF610,ZNF718,ZNF728,ZNF90,ZNF506,DOCK5,GFM2,IDH1,NELL2,RAB21,ZNF141 |
| GO:MF | RNA polymerase II transcription regulatory region sequence-specific DNA binding | GO:0000977 | 3.88981E-05 | 4.410071776 | 1387 | 147 | 30 | 20246 | ZNF736,ZNF257,CTCF,ZNF66,KDM6A,ZNF431,ZNF721,ZNF726,ZNF729,ZNF761,ZNF845,ZNF93,MTF2,ZNF708,ZNF595,ZNF765,ZNF98,ZFHX4,ZNF91,ZNF860,ZNF737,PAX8,ZNF429,ZNF610,ZNF718,ZNF728,ZNF90,ZNF506,HSF5,ZNF141 |
| GO:MF | cation binding | GO:0043169 | 7.95059E-05 | 4.099600522 | 4495 | 147 | 61 | 20246 | TTN,KMT2C,ZNF736,ZNF257,HMCN1,RNF213,CTCF,USP32,TYW1,ITGA2,ZNF66,KDM6A,ZNF431,PCLO,ZNF721,CACNA1H,FLT4,RYR1,ZNF726,ZNF729,RYR3,ZNF761,ZNF845,ZNF93,CACNA1A,CELSR2,CELSR3,MTF2,NF1,UTRN,ZMIZ2,ZNF708,ZNF595,ZNF765,ZNF98,CELSR1,DGKK,EHD2,GTF2E1,PLA2G4C,PRKCG,SLC25A23,ZFHX4,ZNF830,ZNF91,ZNF860,ZNF737,ADAMTS10,COLEC12,CPXM1,EGFLAM,METAP1,ZNF429,ZNF610,ZNF718,ZNF728,ZNF90,ZNF506,IDH1,NELL2,ZNF141 |
| GO:MF | metal ion binding | GO:0046872 | 8.8553E-05 | 4.052796725 | 4396 | 147 | 60 | 20246 | TTN,KMT2C,ZNF736,ZNF257,HMCN1,RNF213,CTCF,USP32,TYW1,ITGA2,ZNF66,KDM6A,ZNF431,PCLO,ZNF721,CACNA1H,FLT4,RYR1,ZNF726,ZNF729,RYR3,ZNF761,ZNF845,ZNF93,CACNA1A,CELSR2,CELSR3,MTF2,UTRN,ZMIZ2,ZNF708,ZNF595,ZNF765,ZNF98,CELSR1,DGKK,EHD2,GTF2E1,PLA2G4C,PRKCG,SLC25A23,ZFHX4,ZNF830,ZNF91,ZNF860,ZNF737,ADAMTS10,COLEC12,CPXM1,EGFLAM,METAP1,ZNF429,ZNF610,ZNF718,ZNF728,ZNF90,ZNF506,IDH1,NELL2,ZNF141 |
| GO:MF | double-stranded DNA binding | GO:0003690 | 0.000145508 | 3.837113737 | 1644 | 147 | 32 | 20246 | ZNF736,ZNF257,CTCF,PRKDC,ZNF66,KDM6A,ZNF431,ZNF721,ZNF726,ZNF729,ZNF761,ZNF845,ZNF93,MTF2,ZNF708,ZNF595,ZNF765,ZNF98,ZFHX4,ZNF91,ZNF860,ZNF737,LSM14A,PAX8,ZNF429,ZNF610,ZNF718,ZNF728,ZNF90,ZNF506,HSF5,ZNF141 |
| GO:MF | transcription cis-regulatory region binding | GO:0000976 | 0.000171939 | 3.764624804 | 1485 | 147 | 30 | 20246 | ZNF736,ZNF257,CTCF,ZNF66,KDM6A,ZNF431,ZNF721,ZNF726,ZNF729,ZNF761,ZNF845,ZNF93,MTF2,ZNF708,ZNF595,ZNF765,ZNF98,ZFHX4,ZNF91,ZNF860,ZNF737,PAX8,ZNF429,ZNF610,ZNF718,ZNF728,ZNF90,ZNF506,HSF5,ZNF141 |
| GO:MF | transcription regulatory region nucleic acid binding | GO:0001067 | 0.000174439 | 3.758355604 | 1486 | 147 | 30 | 20246 | ZNF736,ZNF257,CTCF,ZNF66,KDM6A,ZNF431,ZNF721,ZNF726,ZNF729,ZNF761,ZNF845,ZNF93,MTF2,ZNF708,ZNF595,ZNF765,ZNF98,ZFHX4,ZNF91,ZNF860,ZNF737,PAX8,ZNF429,ZNF610,ZNF718,ZNF728,ZNF90,ZNF506,HSF5,ZNF141 |
| GO:MF | DNA-binding transcription factor activity | GO:0003700 | 0.000339522 | 3.469131801 | 1448 | 147 | 29 | 20246 | ZNF736,ZNF257,CTCF,ZNF66,ZNF431,ZNF721,ZNF726,ZNF729,ZNF761,ZNF845,ZNF93,ZNF708,ZNF595,ZNF765,ZNF98,ZFHX4,ZNF91,ZNF860,ZNF737,PAX8,SKIDA1,ZNF429,ZNF610,ZNF718,ZNF728,ZNF90,ZNF506,HSF5,ZNF141 |
| GO:MF | sequence-specific double-stranded DNA binding | GO:1990837 | 0.000410015 | 3.387199879 | 1547 | 147 | 30 | 20246 | ZNF736,ZNF257,CTCF,ZNF66,KDM6A,ZNF431,ZNF721,ZNF726,ZNF729,ZNF761,ZNF845,ZNF93,MTF2,ZNF708,ZNF595,ZNF765,ZNF98,ZFHX4,ZNF91,ZNF860,ZNF737,PAX8,ZNF429,ZNF610,ZNF718,ZNF728,ZNF90,ZNF506,HSF5,ZNF141 |
| GO:MF | microtubule motor activity | GO:0003777 | 0.000422593 | 3.37407741 | 68 | 147 | 7 | 20246 | KIF14,DNAH11,DYNC2H1,DNAH3,DNAH5,KIFC2,KIF7 |
| GO:MF | sequence-specific DNA binding | GO:0043565 | 0.000554106 | 3.256407309 | 1657 | 147 | 31 | 20246 | ZNF736,ZNF257,CTCF,ZNF66,KDM6A,ZNF431,ZNF721,ZNF726,ZNF729,ZNF761,ZNF845,ZNF93,MTF2,ZNF708,ZNF595,ZNF765,ZNF98,GOLGB1,ZFHX4,ZNF91,ZNF860,ZNF737,PAX8,ZNF429,ZNF610,ZNF718,ZNF728,ZNF90,ZNF506,HSF5,ZNF141 |
| GO:MF | DNA binding | GO:0003677 | 0.000882304 | 3.054381678 | 2542 | 147 | 40 | 20246 | KMT2C,ZNF736,ZNF257,CHD9,CTCF,PRKDC,ZNF66,KDM6A,ZNF431,CHD8,ZNF721,ZNF726,ZNF729,ZNF761,ZNF845,ZNF93,MTF2,ZNF708,ZNF595,ZNF765,ZNF98,GOLGB1,PARP4,POLDIP2,UBN1,ZFHX4,ZNF91,ZNF860,ZNF737,BTAF1,LSM14A,PAX8,ZNF429,ZNF610,ZNF718,ZNF728,ZNF90,ZNF506,HSF5,ZNF141 |
| GO:MF | DNA-binding transcription factor activity, RNA polymerase II-specific | GO:0000981 | 0.001068593 | 2.971187684 | 1360 | 147 | 27 | 20246 | ZNF736,ZNF257,CTCF,ZNF66,ZNF431,ZNF721,ZNF726,ZNF729,ZNF761,ZNF845,ZNF93,ZNF708,ZNF595,ZNF765,ZNF98,ZFHX4,ZNF91,ZNF860,ZNF737,PAX8,ZNF429,ZNF610,ZNF718,ZNF728,ZNF90,ZNF506,ZNF141 |
| GO:MF | transcription regulator activity | GO:0140110 | 0.002189067 | 2.659741005 | 1954 | 147 | 33 | 20246 | KMT2C,ZNF736,ZNF257,CTCF,ZNF66,ZNF431,CTNNB1,ZNF721,ZNF726,ZNF729,ZNF761,ZNF845,ZNF93,ZNF708,ZNF595,ZNF765,ZNF98,TRRAP,ZFHX4,ZNF91,ZNF860,ZNF737,BTAF1,PAX8,SKIDA1,ZNF429,ZNF610,ZNF718,ZNF728,ZNF90,ZNF506,HSF5,ZNF141 |
| GO:MF | minus-end-directed microtubule motor activity | GO:0008569 | 0.004040508 | 2.393564071 | 17 | 147 | 4 | 20246 | DNAH11,DYNC2H1,DNAH3,DNAH5 |
| GO:MF | cytoskeletal motor activity | GO:0003774 | 0.014377904 | 1.842304426 | 115 | 147 | 7 | 20246 | KIF14,DNAH11,DYNC2H1,DNAH3,DNAH5,KIFC2,KIF7 |
| GO:MF | polypeptide conformation or assembly isomerase activity | GO:0120544 | 0.019983722 | 1.699323624 | 121 | 147 | 7 | 20246 | KIF14,DNAH11,DYNC2H1,DNAH3,DNAH5,KIFC2,KIF7 |
| GO:MF | dynein light intermediate chain binding | GO:0051959 | 0.032666626 | 1.485895724 | 28 | 147 | 4 | 20246 | DNAH11,DYNC2H1,DNAH3,DNAH5 |
| GO:MF | macromolecular conformation isomerase activity | GO:0120543 | 0.036063146 | 1.44293639 | 290 | 147 | 10 | 20246 | CHD9,KIF14,DNAH11,CHD8,DYNC2H1,DNAH3,DNAH5,KIFC2,KIF7,BTAF1 |
| GO:BP | regulation of cellular process | GO:0050794 | 0.000476501 | 3.321936297 | 11946 | 144 | 111 | 21026 | TTN,APC,KMT2C,ZNF736,ZNF257,CDC27,RNF213,NEB,KIF14,CTCF,TOGARAM1,USP32,ITGA2,PRKDC,RXFP2,ZNF66,KDM6A,ZNF431,DNAH11,PCLO,CHD8,CTNNB1,ZNF721,FLT4,RYR1,SIPA1L2,ZNF726,ZNF729,DYNC2H1,RYR3,ZNF761,ZNF845,ZNF93,CACNA1A,CELSR2,CELSR3,FAM83H,GNL1,MTF2,NF1,PCM1,SASH1,SZT2,TNXB,UBE3B,UTRN,ZMIZ2,ZNF708,DOCK1,ZNF595,ZNF765,ZNF98,ANK3,CASKIN1,CELSR1,DGKK,EHD2,ELP2,FCRL3,GOLGB1,IP6K2,KIF7,MST1R,PLA2G4C,POLDIP2,PRKCG,RIN2,SLC25A23,TRRAP,ZFHX4,ZNF830,ZNF91,ZNF860,ZNF737,ADGRA3,BRD3,BTAF1,CKAP5,COLEC12,CSMD3,CYFIP2,EGFLAM,FRY,HOMER1,KCNJ9,LSM14A,MACC1,METAP1,NUMB,PAX8,PTCRA,RPL10,SKIDA1,SPAG5,ZNF429,ZNF610,ZNF718,ZNF728,ZNF90,ZNF506,DOCK5,COL4A3,F3,HSF5,IBTK,IDH1,IVNS1ABP,METTL8,RAB21,WDR75,ZNF141 |
| GO:BP | regulation of biological process | GO:0050789 | 0.000665434 | 3.176895271 | 12336 | 144 | 113 | 21026 | TTN,APC,KMT2C,ZNF736,ZNF257,CDC27,RNF213,NEB,KIF14,CTCF,TOGARAM1,USP32,ITGA2,PRKDC,RXFP2,ZNF66,KDM6A,ZNF431,DNAH11,PCLO,CHD8,CTNNB1,ZNF721,CACNA1H,FLT4,RYR1,SIPA1L2,ZNF726,ZNF729,DYNC2H1,RYR3,ZNF761,ZNF845,ZNF93,CACNA1A,CELSR2,CELSR3,FAM83H,GNL1,MTF2,NF1,PCM1,SASH1,SZT2,TNXB,UBE3B,UTRN,ZMIZ2,ZNF708,DOCK1,ZNF595,ZNF765,ZNF98,ANK3,CASKIN1,CELSR1,DGKK,EHD2,ELP2,FCRL3,GOLGB1,IP6K2,KIF7,MST1R,PLA2G4C,POLDIP2,PRKCG,RIN2,SLC25A23,TRRAP,ZFHX4,ZNF830,ZNF91,ZNF860,ZNF737,ADGRA3,BRD3,BTAF1,CKAP5,COLEC12,CSMD3,CYFIP2,EGFLAM,FRY,HOMER1,KCNJ9,LSM14A,MACC1,METAP1,NLRP4,NUMB,PAX8,PTCRA,RPL10,SKIDA1,SPAG5,ZNF429,ZNF610,ZNF718,ZNF728,ZNF90,ZNF506,DOCK5,COL4A3,F3,HSF5,IBTK,IDH1,IVNS1ABP,METTL8,RAB21,WDR75,ZNF141 |
| GO:BP | regulation of transcription by RNA polymerase II | GO:0006357 | 0.000673036 | 3.171961976 | 2578 | 144 | 40 | 21026 | KMT2C,ZNF736,ZNF257,CTCF,PRKDC,ZNF66,KDM6A,ZNF431,CHD8,CTNNB1,ZNF721,ZNF726,ZNF729,ZNF761,ZNF845,ZNF93,ZMIZ2,ZNF708,ZNF595,ZNF765,ZNF98,ELP2,TRRAP,ZFHX4,ZNF91,ZNF860,ZNF737,BRD3,MACC1,PAX8,RPL10,SKIDA1,ZNF429,ZNF610,ZNF718,ZNF728,ZNF90,ZNF506,HSF5,ZNF141 |
| GO:BP | transcription by RNA polymerase II | GO:0006366 | 0.000905311 | 3.043202367 | 2711 | 144 | 41 | 21026 | KMT2C,ZNF736,ZNF257,CTCF,PRKDC,ZNF66,KDM6A,ZNF431,CHD8,CTNNB1,ZNF721,ZNF726,ZNF729,ZNF761,ZNF845,ZNF93,ZMIZ2,ZNF708,ZNF595,ZNF765,ZNF98,ELP2,GTF2E1,TRRAP,ZFHX4,ZNF91,ZNF860,ZNF737,BRD3,MACC1,PAX8,RPL10,SKIDA1,ZNF429,ZNF610,ZNF718,ZNF728,ZNF90,ZNF506,HSF5,ZNF141 |
| GO:BP | regulation of primary metabolic process | GO:0080090 | 0.002904538 | 2.536922942 | 5390 | 144 | 63 | 21026 | APC,KMT2C,ZNF736,ZNF257,RNF213,KIF14,CTCF,ITGA2,PRKDC,ZNF66,KDM6A,ZNF431,CHD8,CTNNB1,ZNF721,FLT4,ZNF726,ZNF729,ZNF761,ZNF845,ZNF93,CELSR2,MTF2,NF1,SASH1,ZMIZ2,ZNF708,ZNF595,ZNF765,ZNF98,ELP2,GOLGB1,IP6K2,MST1R,PRKCG,TRRAP,ZFHX4,ZNF91,ZNF860,ZNF737,BRD3,BTAF1,CYFIP2,FRY,LSM14A,MACC1,METAP1,PAX8,RPL10,SKIDA1,ZNF429,ZNF610,ZNF718,ZNF728,ZNF90,ZNF506,HSF5,IBTK,IDH1,IVNS1ABP,METTL8,WDR75,ZNF141 |
| GO:BP | DNA-templated transcription | GO:0006351 | 0.004656675 | 2.331924086 | 3549 | 144 | 47 | 21026 | KMT2C,ZNF736,ZNF257,CTCF,PRKDC,ZNF66,KDM6A,ZNF431,CHD8,CTNNB1,ZNF721,ZNF726,ZNF729,ZNF761,ZNF845,ZNF93,CELSR2,MTF2,ZMIZ2,ZNF708,ZNF595,ZNF765,ZNF98,ELP2,GOLGB1,GTF2E1,TRRAP,ZFHX4,ZNF91,ZNF860,ZNF737,BRD3,BTAF1,MACC1,PAX8,RPL10,SKIDA1,ZNF429,ZNF610,ZNF718,ZNF728,ZNF90,ZNF506,HSF5,IVNS1ABP,WDR75,ZNF141 |
| GO:BP | biological regulation | GO:0065007 | 0.006279441 | 2.202079031 | 12743 | 144 | 113 | 21026 | TTN,APC,KMT2C,ZNF736,ZNF257,CDC27,RNF213,NEB,KIF14,CTCF,TOGARAM1,USP32,ITGA2,PRKDC,RXFP2,ZNF66,KDM6A,ZNF431,DNAH11,PCLO,CHD8,CTNNB1,ZNF721,CACNA1H,FLT4,RYR1,SIPA1L2,ZNF726,ZNF729,DYNC2H1,RYR3,ZNF761,ZNF845,ZNF93,CACNA1A,CELSR2,CELSR3,FAM83H,GNL1,MTF2,NF1,PCM1,SASH1,SZT2,TNXB,UBE3B,UTRN,ZMIZ2,ZNF708,DOCK1,ZNF595,ZNF765,ZNF98,ANK3,CASKIN1,CELSR1,DGKK,EHD2,ELP2,FCRL3,GOLGB1,IP6K2,KIF7,MST1R,PLA2G4C,POLDIP2,PRKCG,RIN2,SLC25A23,TRRAP,ZFHX4,ZNF830,ZNF91,ZNF860,ZNF737,ADGRA3,BRD3,BTAF1,CKAP5,COLEC12,CSMD3,CYFIP2,EGFLAM,FRY,HOMER1,KCNJ9,LSM14A,MACC1,METAP1,NLRP4,NUMB,PAX8,PTCRA,RPL10,SKIDA1,SPAG5,ZNF429,ZNF610,ZNF718,ZNF728,ZNF90,ZNF506,DOCK5,COL4A3,F3,HSF5,IBTK,IDH1,IVNS1ABP,METTL8,RAB21,WDR75,ZNF141 |
| GO:BP | regulation of DNA-templated transcription | GO:0006355 | 0.009197341 | 2.036337691 | 3409 | 144 | 45 | 21026 | KMT2C,ZNF736,ZNF257,CTCF,PRKDC,ZNF66,KDM6A,ZNF431,CHD8,CTNNB1,ZNF721,ZNF726,ZNF729,ZNF761,ZNF845,ZNF93,CELSR2,MTF2,ZMIZ2,ZNF708,ZNF595,ZNF765,ZNF98,ELP2,GOLGB1,TRRAP,ZFHX4,ZNF91,ZNF860,ZNF737,BRD3,BTAF1,MACC1,PAX8,RPL10,SKIDA1,ZNF429,ZNF610,ZNF718,ZNF728,ZNF90,ZNF506,HSF5,WDR75,ZNF141 |
| GO:BP | regulation of RNA biosynthetic process | GO:2001141 | 0.010675707 | 1.971603343 | 3428 | 144 | 45 | 21026 | KMT2C,ZNF736,ZNF257,CTCF,PRKDC,ZNF66,KDM6A,ZNF431,CHD8,CTNNB1,ZNF721,ZNF726,ZNF729,ZNF761,ZNF845,ZNF93,CELSR2,MTF2,ZMIZ2,ZNF708,ZNF595,ZNF765,ZNF98,ELP2,GOLGB1,TRRAP,ZFHX4,ZNF91,ZNF860,ZNF737,BRD3,BTAF1,MACC1,PAX8,RPL10,SKIDA1,ZNF429,ZNF610,ZNF718,ZNF728,ZNF90,ZNF506,HSF5,WDR75,ZNF141 |
| GO:CC | supramolecular polymer | GO:0099081 | 8.34294E-05 | 4.078681104 | 1076 | 149 | 24 | 22149 | TTN,APC,NEB,KIF14,TOGARAM1,EML5,DNAH11,CTNNB1,RYR1,DYNC2H1,RYR3,DNAH3,DNAH5,FAM83H,KIFC2,ANK3,KIF7,PARP4,ADAMTS10,CKAP5,FRG1,HOMER1,SPAG5,COL4A3 |
| GO:CC | supramolecular fiber | GO:0099512 | 0.0002827 | 3.548673468 | 1068 | 149 | 23 | 22149 | TTN,APC,NEB,KIF14,TOGARAM1,EML5,DNAH11,CTNNB1,RYR1,DYNC2H1,RYR3,DNAH3,DNAH5,FAM83H,KIFC2,ANK3,KIF7,PARP4,ADAMTS10,CKAP5,FRG1,HOMER1,SPAG5 |
| GO:CC | Z disc | GO:0030018 | 0.001171942 | 2.931094047 | 132 | 149 | 8 | 22149 | TTN,NEB,CTNNB1,RYR1,RYR3,ANK3,FRG1,HOMER1 |
| GO:CC | supramolecular complex | GO:0099080 | 0.001478701 | 2.830119684 | 1448 | 149 | 26 | 22149 | TTN,APC,NEB,KIF14,TOGARAM1,EML5,DNAH11,CTNNB1,RYR1,DYNC2H1,RYR3,DNAH3,DNAH5,FAM83H,KIFC2,PRRC2C,ANK3,KIF7,PARP4,ADAMTS10,CKAP5,FRG1,HOMER1,LSM14A,SPAG5,COL4A3 |
| GO:CC | I band | GO:0031674 | 0.002604946 | 2.584201222 | 147 | 149 | 8 | 22149 | TTN,NEB,CTNNB1,RYR1,RYR3,ANK3,FRG1,HOMER1 |
| GO:CC | cytoskeleton | GO:0005856 | 0.005354035 | 2.271318754 | 2474 | 149 | 35 | 22149 | TTN,APC,CDC27,NEB,KIF14,TOGARAM1,PPP4R3A,EML5,DNAH11,PCLO,MPRIP,CTNNB1,DYNC2H1,DNAH3,DNAH5,FAM83H,KIFC2,PCM1,UTRN,ANK3,EHD2,KIF7,LYST,MST1R,PARP4,POLDIP2,UBN1,CCDC81,CKAP5,FRY,LRRC43,LSM14A,SPAG5,DOCK5,IVNS1ABP |
| GO:CC | microtubule | GO:0005874 | 0.01089427 | 1.962801855 | 492 | 149 | 13 | 22149 | APC,KIF14,TOGARAM1,EML5,DNAH11,DYNC2H1,DNAH3,DNAH5,KIFC2,KIF7,PARP4,CKAP5,SPAG5 |
| GO:CC | microtubule cytoskeleton | GO:0015630 | 0.01184602 | 1.926427542 | 1436 | 149 | 24 | 22149 | APC,CDC27,KIF14,TOGARAM1,PPP4R3A,EML5,DNAH11,CTNNB1,DYNC2H1,DNAH3,DNAH5,KIFC2,PCM1,EHD2,KIF7,LYST,PARP4,POLDIP2,UBN1,CCDC81,CKAP5,FRY,LSM14A,SPAG5 |
| GO:CC | microtubule associated complex | GO:0005875 | 0.038107787 | 1.418986272 | 159 | 149 | 7 | 22149 | KIF14,DNAH11,DYNC2H1,DNAH3,DNAH5,KIFC2,KIF7 |
| GO:CC | sarcomere | GO:0030017 | 0.043526964 | 1.361241627 | 218 | 149 | 8 | 22149 | TTN,NEB,CTNNB1,RYR1,RYR3,ANK3,FRG1,HOMER1 |
| KEGG | Herpes simplex virus 1 infection | KEGG:05168 | 1.51E-09 | 8.822371705 | 506 | 80 | 24 | 8484 | ZNF736,ZNF257,ZNF66,ZNF431,ZNF721,ZNF726,ZNF729,ZNF761,ZNF845,ZNF93,ZNF708,ZNF595,ZNF765,ZNF98,ZNF91,ZNF860,ZNF737,ZNF429,ZNF610,ZNF718,ZNF728,ZNF90,ZNF506,ZNF141 |
| TF | Factor: PBX3; motif: NNNTGAGTGACAGGNNN | TF:M09781 | 0.000970711 | 3.012910158 | 290 | 152 | 13 | 20092 | ZNF257,ZNF66,ZNF431,ZNF726,ZNF93,ZNF98,ZNF91,ZNF737,MACC1,ZNF429,ZNF718,ZNF90,ZNF141 |
| TF | Factor: NF-Y; motif: NCTGATTGGYTASY; match class: 1 | TF:M00209_1 | 0.001077194 | 2.967706187 | 703 | 152 | 20 | 20092 | ZNF736,CTCF,TYW1,ZNF66,ZNF431,ZNF726,ZNF729,ZNF93,SASH1,ZNF595,GOLGB1,ZNF737,GOLGA8H,NLRP4,SKIDA1,ZNF429,ZNF728,ZNF90,ZNF506,ZNF141 |
| TF | Factor: NFYA; motif: RRCCAATCAGN; match class: 1 | TF:M09816_1 | 0.006941909 | 2.15852111 | 292 | 152 | 12 | 20092 | ZNF66,ZNF431,ZNF726,ZNF729,ZNF93,ZNF595,ZNF737,SKIDA1,ZNF429,ZNF728,ZNF506,ZNF141 |
| TF | Factor: PREP-1; motif: TGANTGRCAGS; match class: 1 | TF:M10032_1 | 0.021436786 | 1.668840326 | 94 | 152 | 7 | 20092 | TTN,ZNF66,ZNF431,ZNF726,ZNF91,ZNF718,ZNF141 |
| TF | Factor: PKNOX1; motif: NNTGAGTGACAGNNN | TF:M12559 | 0.023855728 | 1.622407331 | 448 | 152 | 14 | 20092 | TTN,ZNF257,ZNF66,ZNF431,ZNF726,ZNF93,ZNF98,ZNF91,ZNF737,MACC1,ZNF429,ZNF718,ZNF90,ZNF141 |
| MIRNA | hsa-mir-4504 | MIRNA:hsa-mir-4504 | 0.001248544 | 2.903596101 | 816 | 141 | 22 | 16638 | ZNF257,RNF213,CHD9,CTCF,ZNF431,MPRIP,ZNF761,CELSR2,FAM83H,GNL1,UBE3B,ZNF708,ZNF765,ZNF91,BRD3,BTAF1,LSM14A,PPM1H,ZNF718,ZNF506,U2SURP,RAB21 |
| MIRNA | hsa-mir-4768-5p | MIRNA:hsa-mir-4768-5p | 0.001390608 | 2.856795253 | 1787 | 141 | 35 | 16638 | ZNF257,RNF213,CHD9,USP32,ITGA2,ZNF431,MPRIP,CTNNB1,ZNF721,ZNF761,ZNF845,GNL1,MTF2,SASH1,SZT2,TNXB,UTRN,ZNF708,ZNF765,LYST,ZNF91,ZNF737,CYFIP2,EGFLAM,FRY,PAX8,PPM1H,ZNF718,ZNF90,ZNF506,DOCK5,GFM2,HSF5,IBTK,METTL8 |
| MIRNA | hsa-mir-548an | MIRNA:hsa-mir-548an | 0.011411118 | 1.942671786 | 1467 | 141 | 29 | 16638 | ZNF736,ZNF257,RNF213,KIF14,NBPF12,ZNF431,CTNNB1,ZNF721,ZNF761,ZNF845,ZNF93,FAM83H,NBPF15,PRRC2C,SASH1,SZT2,UBE3B,ZNF708,DOCK1,ZNF98,IP6K2,RIN2,ZFHX4,ZNF91,ZNF737,ZNF90,ZNF506,IVNS1ABP,RAB21 |
| MIRNA | hsa-mir-549a | MIRNA:hsa-mir-549a | 0.027003565 | 1.568578891 | 43 | 141 | 5 | 16638 | NBPF12,MPRIP,GNL1,NBPF15,ZNF708 |
| MIRNA | hsa-mir-4484 | MIRNA:hsa-mir-4484 | 0.043397325 | 1.362537043 | 599 | 141 | 16 | 16638 | ZNF736,ZNF257,KIF14,ZNF431,ZNF721,ZNF761,ZNF845,ZNF93,NF1,ZNF708,DOCK1,ZNF98,ZNF91,ZNF737,ZNF90,ZNF506 |
| MIRNA | hsa-mir-3674 | MIRNA:hsa-mir-3674 | 0.043654986 | 1.359966148 | 804 | 141 | 19 | 16638 | ZNF257,RNF213,PRKDC,ZNF431,ZNF721,ZNF761,ZNF93,NBPF15,SASH1,ZNF708,ANK3,ZNF91,ZNF737,CSMD3,HOMER1,LSM14A,ZNF506,RAB21,ZNF141 |

**Supplementary Table S4**: *Pathway analysis of genes mutated in at least 30% of all ARLD tumours; patients with multifocal disease are only represented once.*

| source | term_name | term_id | adjusted_p_value | negative_log10_of_adjusted_p_value | term_size | query_size | intersection_size | effective_domain_size | intersections |
|---|---|---|---|---|---|---|---|---|---|
| GO:BP | lipoprotein transport | GO:0042953 | 0.044773557 | 1.348978403 | 20 | 10 | 2 | 21026 | MIA2,LRP1 |
| GO:BP | lipoprotein localization | GO:0044872 | 0.049474008 | 1.305622902 | 21 | 10 | 2 | 21026 | MIA2,LRP1 |
| WP | Regulation of Wnt B catenin signaling by small molecule compounds | WP:WP3664 | 0.000694656 | 3.15823022 | 17 | 3 | 2 | 8752 | CTNNB1,LRP1 |
| CORUM | TCF4-CTNNB1 complex | CORUM:5262 | 0.049987986 | 1.301134359 | 1 | 3 | 1 | 3383 | CTNNB1 |
| CORUM | Cell-cell junction complex (CDH1-CTNNB1) | CORUM:5281 | 0.049987986 | 1.301134359 | 1 | 3 | 1 | 3383 | CTNNB1 |
| CORUM | CTNNB1-DNMT1 complex | CORUM:6336 | 0.049987986 | 1.301134359 | 1 | 3 | 1 | 3383 | CTNNB1 |
| CORUM | CD44-LRP1 complex | CORUM:7535 | 0.049987986 | 1.301134359 | 1 | 3 | 1 | 3383 | LRP1 |

**Supplementary Table S5**: *Pathway analysis of genes mutated in at least 30% of all HCV-related tumours; patients with multifocal disease are only represented once.*

| term_id | adjusted_p_value | negative_log10_of_adjusted_p_value | term_size | query_size | intersection_size | effective_domain_size | intersections |
|---|---|---|---|---|---|---|---|
| GO:0008569 | 0.000681778 | 3.166356817 | 17 | 29 | 3 | 20246 | DNAH11,DNAH1,DYNC1H1 |
| GO:0051959 | 0.003249947 | 2.488123763 | 28 | 29 | 3 | 20246 | DNAH11,DNAH1,DYNC1H1 |
| GO:0051015 | 0.00417151 | 2.379706715 | 209 | 29 | 5 | 20246 | TTN,SPTBN4,MPRIP,SPTAN1,SYNE1 |
| GO:0005198 | 0.005979805 | 2.223312977 | 1106 | 29 | 9 | 20246 | TTN,DST,MUC17,PCLO,SPTBN4,EPPK1,SPTAN1,AHNAK,LAMA2 |
| GO:0045505 | 0.007641685 | 2.116810895 | 37 | 29 | 3 | 20246 | DNAH11,DNAH1,DYNC1H1 |
| GO:0003779 | 0.012483681 | 1.903657326 | 442 | 29 | 6 | 20246 | TTN,DST,SPTBN4,MPRIP,SPTAN1,SYNE1 |
| GO:0003777 | 0.047839363 | 1.320214616 | 68 | 29 | 3 | 20246 | DNAH11,DNAH1,DYNC1H1 |
| GO:0097493 | 0.049843398 | 1.302392362 | 12 | 29 | 2 | 20246 | TTN,AHNAK |
| GO:0006996 | 0.005617786 | 2.250434813 | 3594 | 26 | 15 | 21026 | TTN,SMG1,CDC27,DST,PCLO,SPTBN4,DNAH1,DYNC1H1,EPPK1,GCN1,MKI67,SPTAN1,ESPL1,SYNE1,UBR4 |
| GO:0005856 | 9.54E-07 | 6.020349571 | 2474 | 28 | 16 | 22149 | TTN,CDC27,DST,DNAH11,PCLO,SPTBN4,MPRIP,DNAH1,DYNC1H1,EPPK1,SPTAN1,ESPL1,HERC2,SYNE1,AHNAK,UBR4 |
| GO:0015629 | 0.00072638 | 3.138835938 | 534 | 28 | 7 | 22149 | TTN,DST,SPTBN4,MPRIP,SPTAN1,SYNE1,AHNAK |
| GO:0099512 | 0.000857404 | 3.066814339 | 1068 | 28 | 9 | 22149 | TTN,DST,DNAH11,SPTBN4,DNAH1,DYNC1H1,EPPK1,SYNE1,AHNAK |
| GO:0099081 | 0.000911348 | 3.040315948 | 1076 | 28 | 9 | 22149 | TTN,DST,DNAH11,SPTBN4,DNAH1,DYNC1H1,EPPK1,SYNE1,AHNAK |
| GO:0099080 | 0.001277722 | 2.893563792 | 1448 | 28 | 10 | 22149 | TTN,SMG1,DST,DNAH11,SPTBN4,DNAH1,DYNC1H1,EPPK1,SYNE1,AHNAK |
| GO:0043232 | 0.00162891 | 2.788102909 | 6669 | 28 | 20 | 22149 | TTN,SMG1,CDC27,DST,DNAH11,PCLO,SPTBN4,CHD8,MPRIP,DNAH1,DYNC1H1,EPPK1,GCN1,MKI67,SPTAN1,ESPL1,HERC2,SYNE1,AHNAK,UBR4 |
| GO:0043228 | 0.001633029 | 2.787006176 | 6670 | 28 | 20 | 22149 | TTN,SMG1,CDC27,DST,DNAH11,PCLO,SPTBN4,CHD8,MPRIP,DNAH1,DYNC1H1,EPPK1,GCN1,MKI67,SPTAN1,ESPL1,HERC2,SYNE1,AHNAK,UBR4 |
| GO:0042995 | 0.003645582 | 2.438233176 | 2441 | 28 | 12 | 22149 | LRP2,DST,DNAH11,PCLO,SPTBN4,DNAH1,DYNC1H1,EPPK1,SPTAN1,SYNE1,LAMA2,SACS |
| GO:0030056 | 0.006744109 | 2.171075393 | 7 | 28 | 2 | 22149 | DST,EPPK1 |
| GO:0099568 | 0.008593653 | 2.06582218 | 313 | 28 | 5 | 22149 | DST,DNAH11,PCLO,DNAH1,DYNC1H1 |
| GO:0070852 | 0.00898511 | 2.046476581 | 8 | 28 | 2 | 22149 | SPTBN4,SACS |
| GO:0030286 | 0.008991551 | 2.046165373 | 54 | 28 | 3 | 22149 | DNAH11,DNAH1,DYNC1H1 |
| GO:0015630 | 0.009237711 | 2.034435621 | 1436 | 28 | 9 | 22149 | CDC27,DST,DNAH11,DNAH1,DYNC1H1,SPTAN1,ESPL1,HERC2,UBR4 |
| GO:0005938 | 0.010734998 | 1.96919805 | 328 | 28 | 5 | 22149 | DST,PCLO,SPTBN4,DYNC1H1,SPTAN1 |
| GO:0008091 | 0.011543247 | 1.937672014 | 9 | 28 | 2 | 22149 | SPTBN4,SPTAN1 |
| GO:0030424 | 0.030543432 | 1.515082167 | 659 | 28 | 6 | 22149 | LRP2,DST,PCLO,SPTBN4,DYNC1H1,SACS |
| GO:0043005 | 0.041475525 | 1.382208106 | 1355 | 28 | 8 | 22149 | LRP2,DST,PCLO,SPTBN4,DYNC1H1,SYNE1,LAMA2,SACS |
| GO:0030016 | 0.04693789 | 1.328476438 | 240 | 28 | 4 | 22149 | TTN,DST,SYNE1,AHNAK |
| REAC:R-HSA-5083625 | 0.00129038 | 2.889282425 | 16 | 22 | 3 | 11004 | MUC16,MUC17,MUC5B |
| REAC:R-HSA-5083636 | 0.00129038 | 2.889282425 | 16 | 22 | 3 | 11004 | MUC16,MUC17,MUC5B |
| REAC:R-HSA-5083632 | 0.001564861 | 2.805524359 | 17 | 22 | 3 | 11004 | MUC16,MUC17,MUC5B |
| REAC:R-HSA-977068 | 0.004043976 | 2.393191457 | 23 | 22 | 3 | 11004 | MUC16,MUC17,MUC5B |
| REAC:R-HSA-5621480 | 0.005913915 | 2.228124932 | 26 | 22 | 3 | 11004 | MUC16,MUC17,MUC5B |
| HP:0008994 | 0.004511284 | 2.345699821 | 73 | 17 | 5 | 5080 | TTN,DYNC1H1,SPTAN1,SYNE1,LAMA2 |
| HP:0001437 | 0.02476841 | 1.606101867 | 103 | 17 | 5 | 5080 | TTN,DYNC1H1,SYNE1,LAMA2,SACS |
| HP:0002363 | 0.030363091 | 1.517654014 | 285 | 17 | 7 | 5080 | PCLO,DYNC1H1,SPTAN1,SYNE1,DEPDC5,LAMA2,SACS |
| HP:0001319 | 0.041405804 | 1.382938776 | 196 | 17 | 6 | 5080 | DST,PCLO,SPTBN4,HERC2,SYNE1,LAMA2 |

**Supplementary table S6:** *Pathway analysis of genes mutated in at least 30% of all NAFLD-related tumours; patients with multifocal disease are only represented once.*
